# Shifting baselines increase the risk of misinterpreting biodiversity trends

**DOI:** 10.1101/2025.05.07.652594

**Authors:** Ariane Dellavalle, Adam J.M. Devenish, Crinan Jarrett, Nathaniel N.D. Annorbah, Augustus Asamoah, Kwame Boafo, Poppy E. Lane, Jake Owen, Alexandra C. Morel, Mark F. Hulme, Andreanna J. Welch, Ken Norris, Fiona J. Sanderson, Joseph A. Tobias

**Author notes:** Corresponding author: Ariane Dellavalle, Joseph A. Tobias. **Statement of authorship:** JAT, AJMD and AD conceived and developed the study, with input from CJ. AJMD, NNDA, AA, and KB collected data. ACM, KN, MFH and FJS provided additional datasets. AD led integration of datasets and ran all analyses with support from AJMD, CJ, PEL and JO. AD wrote the first version of the manuscript and designed all figures, with input from JAT, CJ, AJMD and AJW. All authors contributed to manuscript revisions and gave permission for publication. **Data accessibility statement:** The data that support the findings of this study are openly available in Dryad at https://datadryad.org/xxxx (full link to be added on acceptance). Author (A. Dellavalle), (A. J. M. Devenish), (C. Jarrett), (N. N. D. Annorbah), (A. Asamoah), (K. Boafo), (P. E. Lane), (J. Owen), (A.C. Morel), (M. F. Hulme), (A. J. Welch), (K. Norris), (F. J. Sanderson), (J. A. Tobias).

## Abstract

Ecological studies quantifying the impact of land-use change on biodiversity may be sensitive to the choice of reference points – or baselines – particularly when sampling across human land-use gradients and other space-for-time comparisons. Much depends on whether the chosen baseline has already undergone shifts in species composition because of hunting, habitat loss and degradation. However, few studies have assessed the influence of shifting baselines on estimates of anthropogenic impacts. Using new survey data from five West African land-use gradients, we examine how habitat patch size and structure influences the estimated impact of land-use change on bird species richness and functional diversity. We show that smaller forests have already lost many forest-dependent birds, particularly those with large body size or specialised ecological niches, leading to reduced estimates of biodiversity loss after deforestation. The steepest biodiversity loss was found in mid-sized forests whereas relatively shallow declines were estimated for the most extensive forests – despite their richer taxonomic and functional diversity. In these larger forest blocks, accurate estimates of biodiversity loss may require longer transects extending beyond the biodiversity ‘shadow’ caused by the more extensive spillover of forest species into the surrounding landscape, potentially linked to source-sink dynamics. These findings suggest that biodiversity assessments are highly sensitive to baseline selection and transect design, highlighting the risk of underestimating land-use impacts unless shifting baselines are carefully considered.

## Introduction

Human-induced land-use change is the leading driver of biodiversity loss (Jaureguiberry et al., 2022), particularly in tropical forests, where conversion to agriculture to meet rising food demands is accelerating species declines and altering ecosystem functions (Jarrett et al., 2024; Maas et al., 2016; Mills et al., 2023; Pendrill et al., 2022). Despite these widespread impacts, relatively few long-term datasets track biodiversity change at fixed locations over time (Dornelas et al., 2018). Our understanding of these changes therefore primarily derives from space-for-time comparisons, comparing current biodiversity levels across land-use gradients as a proxy for temporal changes (Cordier et al., 2021; Gray et al., 2016; Millard et al., 2021; Pickett, 1989; Purvis et al., 2018). One limitation of space-for-time comparisons is the condition of the reference point – or baseline – which may be biased by long-term processes driving biodiversity loss, including megafaunal extinctions (Søndergaard et al., 2025) and landscape history (Damgaard, 2019; Lovell et al., 2023; Stouffer et al., 2021). In these cases, the decline of biodiversity over time results in shifting baselines; that is, an altered perception of what constitutes a “natural” or “intact” state (Soga & Gaston, 2018; Søndergaard et al., 2025), with major implications for understanding the impacts of recent land-use change (Figure 1).

**FIGURE 1.**
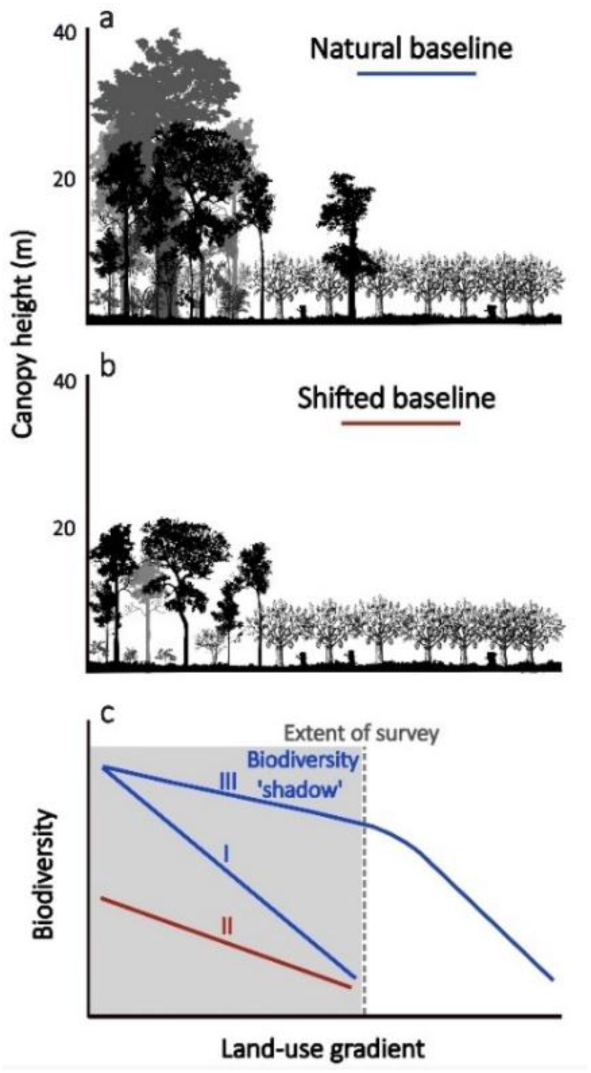
Potential impacts of shifting baselines on estimates of biodiversity loss in tropical forests. Field surveys typically sample biodiversity from forest to agriculture, but the baseline (comparison) sites vary from large, intact, tall-stature forests (**a**) to relatively small, disturbed, low-stature forests (**b**). In addition, post land-use change sites can vary from less intensive (**a**) to more intensive agriculture (**b**). In this context, a ‘natural baseline’ refers to an intact, tall-stature forest, serving as the pre-disturbance reference for biodiversity metrics such as species richness or functional diversity. The choice of baseline in space-for-time comparisons can influence biodiversity loss gradients in two contrasting ways (**c**). First, the slope of biodiversity loss from a natural baseline (I) may be underestimated by a shifted baseline (II) because much biodiversity has already been lost prior to sampling. Alternatively, larger and more intact forests can have a wider biodiversity ‘shadow’ linked to source-sink dynamics, with greater spillover of forest species beyond the boundaries of natural habitat into the surrounding landscape, flattening the gradient of biodiversity change unless survey gradients are lengthened (III).

In many regions, the least-degraded landscapes available as baselines have already experienced significant ecological change. This applies to any study comparing anthropogenic landscapes with relatively small patches of natural habitat. Even when these are fragments of old growth forest, they are likely to have lost many native species owing to species-area relationships (Hanski et al., 2013) and greater exposure to edge effects, reducing the availability of core habitats (Fahrig et al., 2022; Moilanen & Wintle, 2007). These mechanisms, and associated human pressures like hunting, drive non-random local extinctions, disproportionately affecting large-bodied, forest-dependent or dispersal-limited species (Bregman et al., 2014; Peres et al., 2016; Weeks et al., 2023). Species with large body mass or ecological specialisms may be the first to disappear (Bovo et al., 2018; Galetti et al., 2017), driven by resource scarcity and spatial constraints, as well as direct human exploitation, which tends to increase in small fragments (Bregman et al., 2016; Tobias et al., 2013). These species losses mean that taxonomic and functional diversity is already reduced in smaller or more disturbed forests, potentially reducing the slope of biodiversity decline across land-use gradients (Figure 1c II).

Source-sink dynamics and recent landscape history further complicate the interpretation of local-scale biodiversity gradients. In larger forests, source populations are usually larger or more productive, increasing the likelihood of spillover into surrounding degraded or agricultural areas (Brodie et al., 2023; Ewers & Didham, 2006; Lucey et al., 2014), inflating species richness around larger habitat patches and creating a wider ‘biodiversity shadow’. Consequently, over similar land-use gradient lengths, the slope of biodiversity loss may appear flatter around large forest patches compared to shifted baselines (Figure 1c III). These patterns are further nuanced by time lags in species’ responses to forest loss, as extinction debts can temporarily inflate biodiversity around larger forests with more recent clearance around their edges (Brooks et al., 1999), particularly when long-lived species survive in remnants of natural habitat (Daskalova et al., 2020).

The biodiversity impacts of habitat patch size (Kormann et al., 2018; Manu et al., 2007) and land-use change (Bregman et al., 2016; Jarrett et al., 2021) are widely studied and well documented, yet the role of shifting baselines in altering these estimated trends remains poorly understood. A critical question is whether biodiversity gradients calculated from space-for-time comparisons are sensitive to the selection or availability of suitable baselines, and whether such baselines can lead to underestimates of biodiversity loss.

To address this question, we assess the effects of land-use change on bird species richness (SR) and functional diversity (FD) across five West African forests of varying size and landscape history (Figure 2a). We present new or unpublished bird survey data from four study areas in Ghana – Kakum National Park, Ankasa Game Reserve, Dompim, and Ahokwa – and one study area in Sierra Leone – Gola Rainforest National Park. All these areas are remnant patches of a previously continuous forest block, the Upper Guinean rainforest, one of the world’s most fragmented and endangered regions of humid tropical forest (Mittermeier et al., 1999). To compare modern surveys with biodiversity for each forest block, we extracted further bird observation data from literature and online citizen science repositories (e.g. eBird).

**FIGURE 2.**
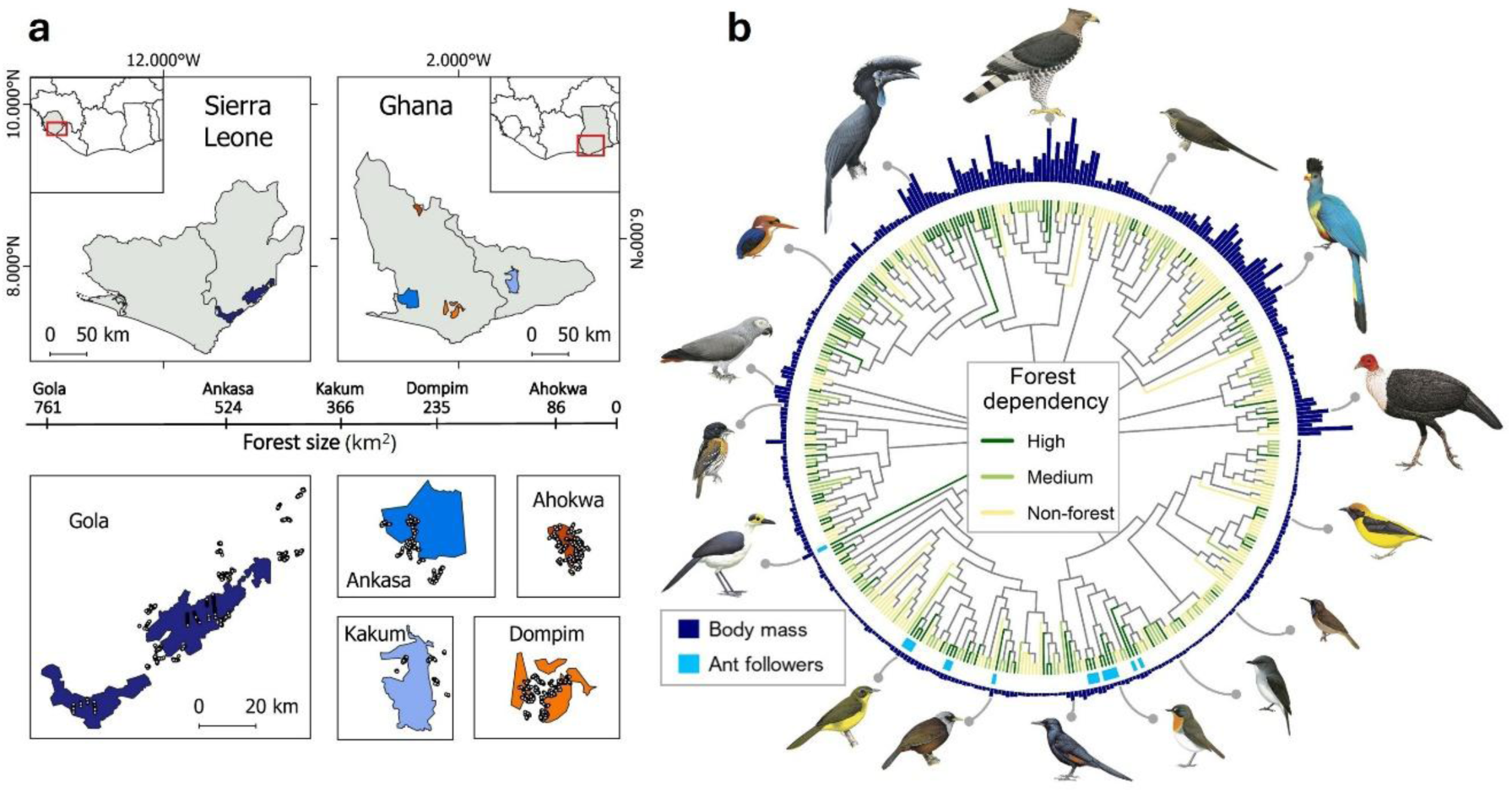
Size, location and avian diversity of sampling sites. In **a**, upper boxes show the geographical distribution of five study areas (Gola, Ankasa, Kakum, Dompim, Ahokwa) in Sierra Leone and Ghana; scale bar across centre shows current variation in land surface area covered by these five forests; lower panels show the distribution of survey transects in each case study (2,479 points across 301 transects). Panel **b** shows the phylogenetic relationship of bird species recorded in the study region (*n* = 413). Variation in body mass is plotted at branch tips of a consensus phylogenetic tree obtained from www.birdtree.org using the Hackett backbone. For ease of visualisation the square root of body mass is shown. Ant followers (i.e. specialised forest birds that follow army ants to prey on invertebrates flushed by the swarm) are highlighted at branch tips. Illustrations show a variety of forest-dependent species, clockwise from top: *Stephanoaetus coronatus*, *Cercococcyx mechowi*, *Corythaeola cristata*, *Agelastes meleagrides*, *Ploceus preussi*, *Cyanomitra cyanolaema*, *Fraseria tessmanni*, *Stiphrornis erythrothorax*, *Onychognathus fulgidus*, *Turdoides atripennis*, *Bleda eximius*, *Picathartes gymnocephalus*, *Smithornis rufolateralis*, *Psittacus erithacus*, *Ispidina lecontei*, *Ceratogymna atrata*.

Bird diversity provides an indicator of ecosystem health (Catterall et al., 2012; Kupsch et al., 2019), with species and trophic guilds varying in their sensitivity to forest degradation (Peters et al., 2008), habitat size (Chasar et al., 2014), and human exploitation (Fingesi et al., 2022). To better understand the role of forest baselines in shaping bird diversity gradients, we classified species into six response groups (invertivores, frugivores, granivores, forest birds, forest birds > 500 g, and ant followers) with well-documented responses to land-use change. Ant followers are ecological specialists that rely on tracking army ants – feeding on invertebrates flushed by ant swarms raiding the forest – and are generally sensitive to forest degradation (Stouffer & Bierregaard, 1995).

Because of recent advances in data availability, birds currently offer the best opportunity to compare diversity gradients across land-use contexts. In particular, comprehensive ecological and morphological trait data linked to key ecological and trophic processes (Pigot et al., 2020) is now available for all bird species (Tobias et al., 2022), providing more detailed insights into diversity gradients, including variation in trophic niches and functional metrics of bird community structure associated with key ecological processes (Bregman et al., 2016; Pigot et al., 2020). We take advantage of this data, in combination with intensive field surveys targeting land-use gradients, to examine the role of baselines in shaping estimated responses of bird diversity to land-use change.

## Methods

### Study sites and survey data

#### Study region and selection of study areas

We surveyed bird communities along forest-to-agriculture gradients in three study areas – Ankasa, Dompim, Ahokwa – in Ghana’s Western Region. The study forests, ranging from 524 km^2^ to 86 km^2^ in size (Figure 2a; see below), are remnant patches of the Upper Guinean rainforest (for details see Supplementary methods). Beyond these forests, the predominant land-use is smallholder farming, with a mosaic of active and abandoned fields. The main crops include maize and cassava, alongside mixed plantations of cocoa, coffee, rubber, and oil palm (Asubonteng et al., 2021).

To expand our dataset, we collated bird survey data collected after 2000 comparing Afrotropical rainforest baselines and surrounding agricultural landscapes. Of seven studies identified, five were removed because of sampling differences (see Data S1). We focused on the remaining two surveys, both conducted within the Upper Guinean rainforest: one in Kakum National Park in Ghana and one in Sierra Leone’s Gola Rainforest National Park, a larger and more intact forest than those remaining in Ghana (Figure 2a).

#### Survey methods

In Ankasa, Dompim and Ahokwa, we conducted point-count surveys (Ankasa; *n* = 444; Dompim: *n* = 403; Ahokwa: *n* = 375) along a land-use gradient using standard methods (Leach et al., 2016; Rurangwa et al., 2022). Surveys in forested habitats extended up to between 2.5 and 6 km from the forest edge towards the forest interior, depending on forest shape and size, while agricultural surveys reached up to 8 km from forests. Birds were surveyed over the course of three field seasons – two in the wet season and one in the dry season – between 2020 and 2021. Surveys were carried out along transects (*n* = 116) consisting of 6-15 survey points at least 200 m apart to minimise double counting (Powell et al., 2015) (for details on sampling design see Supplementary methods). Observations took place between dawn (earliest 04:40) and midday, with each survey lasting 10 minutes. During each count, we recorded all birds seen or heard, noting species, abundance, and whether they were flyovers rather than using the vegetation in the plot (i.e. the circular area of vegetation with 50-m radius, centred on the survey point). Our dataset includes 22,274 records of 231 species.

In both Gola and Kakum, broadly comparable survey approaches were used, with only minor differences in sampling methodology and effort. Both studies employed point-count surveys (Gola: *n* = 817; Kakum: *n* = 440) along land-use gradients, with forest points located between the forest edge and at least 1 km into the forest interior, and agricultural points located up to at least 5 km from the forest. Surveys were conducted over 5–6 field seasons between 2013 and 2018. Surveys were carried out along transects (Gola: *n* = 35; Kakum: *n* = 150), each consisting of 3–26 points spaced at least 200 m apart (for details on sampling design see Supplementary methods). Most surveys took place between 06:15 and 11:30, with a subset in Kakum (*n* = 69) conducted in the late afternoon. Each survey lasted 10 minutes (Kakum) or 15 minutes (Gola), during which all birds seen or heard were identified to species and totalled to give an estimate of relative abundance (number of individuals of each species encountered).

We excluded Gola from analyses assessing bird assemblage responses to land-use change because surveys in its surrounding landscape focused primarily on shaded agriculture under mature tree canopies (Sanderson et al., 2022), whereas agriculture in the Ghanaian landscapes was predominantly unshaded. This key difference in the surrounding matrix, combined with the lack of surveys in low-canopy agricultural areas near Gola (see Figure S1–2), meant that land-use change comparisons with Ghanaian sites would not be ecologically equivalent. However, we retained Gola in analyses investigating the effects of forest size on bird taxonomic and functional diversity within forests. As one of the largest and least disturbed forest sites in the Upper Guinean region, Gola provided a useful baseline for comparison with smaller and more fragmented forests in Ghana.

After integrating these datasets (see Supplementary methods for details of taxonomic matching), our dataset expanded to 46,481 records of 270 species. Each survey was categorised as occurring in either the wet or dry season, following regional monsoonal rainfall patterns (Sanderson et al., 2022; Atiah et al., 2020). To minimise identification errors, all species records were cross-checked against locality data (see Supplementary methods).

Treating each point count as a separate survey can create issues for our analyses due to inadequate representation of the full species assemblage and proximity of neighbouring survey points. To enhance sampling depth and reduce the risk of pseudo-replication, we grouped three point-count sites – located within the same case study and land-use type, and situated no more than 600 m apart from the first to the last point – into a single unit (assemblage surveys, with the bird communities recorded within them hereafter referred to as assemblages). In a small number of cases (*n* = 174), four survey points were grouped together due to transect design constraints (e.g. when a ten-point transect was divided into groups of 3, 3, and 4). This distance exceeds the home range size of most tropical forest birds (Powell et al., 2015) and was chosen to ensure consistency across datasets. We removed species-poor assemblages surveyed at midday (*n* = 3) because the unfavourable time of day for vocalisation likely led to much of the diversity remaining undetected. The final dataset contained 761 assemblages surveyed across forest-to-agriculture gradients.

#### Assessing survey completeness

All field surveys miss species (false negatives), a problem that is particularly pronounced in tropical rainforests, where many species are rare or inconspicuous (Robinson et al., 2018). Additionally, even if we were able to detect all species currently present, some have already gone extinct, highlighting the importance of comparing current observations to historical baselines. To assess our results in this context, and compare our surveys to known and historically recorded regional diversity, we compiled species lists for all our sites from a variety of sources, including protected area checklists (Lepage, 2024), published reports (Dowsett-Lemaire & Dowsett, 2011a, 2011b; Klop et al., 2010), and eBird (Sullivan et al., 2009). These data were available for three study areas – Gola, Ankasa, and Kakum – all of which are national parks frequently visited by birdwatchers. Observations between 2000 and 2024 were extracted from eBird hotspots inside the forest boundaries of Gola Rainforest National Park (*n* = 2), Ankasa Game Reserve (*n* = 4) and Kakum National Park (*n* = 6). Records of species with no credible evidence of likely occurrence within the key protected areas (e.g. species outside their usual range or habitat) were removed. After integrating these records, our dataset comprised 413 species (Figure 2b). For each forest, we compared the number of species detected in our surveys to those not detected, to evaluate how well our surveys captured known and historically recorded bird diversity at each forest block.

### Species traits

Birds respond differently to land-use change in tropical systems depending on their diet and ecological specialisation (Bregman et al., 2014; Jarrett et al., 2021). Some avian functional traits have also been linked to greater sensitivity to land-use change, such as a higher body mass (Gaston & Blackburn, 1995) or lower Hand-wing Index (Weeks et al., 2023). Therefore, we obtained habitat, diet, and morphological functional traits for all 413 species in our dataset (Figure 2b) to model the effects of forest baseline and land-use change separately for different response groups. Species could belong to more than one response group (e.g. invertivore, forest bird, and ant follower; see Data S1).

We compiled forest dependency data from BirdLife International (2024), classifying species into high, medium, low or no forest dependency. In some cases, these forest dependency data were poorly aligned with our study system, for example because of regional differences in habitat use by particular species, or because occasional sightings of forest-based species outside forest habitats have led to them being classified as habitat generalists. We reviewed and amended classifications based on literature, citizen science data (e.g. www.ebird.org) and expert knowledge from the field. When species only occur in non-forest habitats infrequently or in a narrow geographical area, we classified them as high forest dependency for the purposes of our study (for details see Supplementary methods, and Data S1). We use the term ‘forest birds’ exclusively for species with high forest dependency. To account for variation in sensitivity to habitat disturbance and spatial constraints in smaller forest patches, we identified a subset of forest birds as ant followers because this ecological specialism is associated with high sensitivity to environmental change (Martínez et al., 2021; Stouffer & Bierregaard, 1995). In a West African context, ant followers usually follow *Dorylus* army ants, and the distinction between obligate and facultative ant followers is not as well researched as in the Neotropics (Martínez et al., 2021). We used published information (e.g. Billerman et al., 2022; Peters et al., 2008; Waltert et al., 2024) to identify study species that routinely or frequently follow army ant swarms, discounting species which are occasionally observed at swarms (for details see Supplementary methods).

Through their trophic interactions, birds play a crucial role in key ecological processes and services, such as seed dispersal by frugivores and predation of pests and herbivorous insects by invertivores. We extracted trophic niche data from the AVONET database (Tobias et al., 2022) which divides species into nine trophic niches: aquatic predator, frugivore, granivore, herbivore (aquatic), herbivore (terrestrial), invertivore, nectarivore, scavenger, and vertivore. Species are classified into trophic niches based on which food resource they obtain at least 60% of their diet from. Species which exploit a range of food resources similarly are classed as omnivores.

To calculate functional diversity of assemblages we compiled species-mean morphological traits from AVONET (Tobias et al., 2022) for eight biometric traits: beak length (culmen), beak length (tip-to-nares distance), beak depth, beak width, tail length, tarsus length, Hand-wing Index (a measure of wing elongation) and wing length. We also extracted body mass data for each species to identify large forest birds, which are likely to be particularly vulnerable to land-use change due to large home ranges and which have been shown to mediate key ecological services, including seed dispersal over long distances (Bovo et al., 2018; Gaston & Blackburn, 1995). Average values for all morphometric traits were derived from measurements of approximately 8 individuals (mean = 8.1; SD = 14.9) per species (Tobias et al., 2022).

To present our data in the context of evolutionary relationships, we generated a consensus phylogeny for the 413 species in our dataset based on a sample of 100 trees obtained from the most recent global bird phylogeny (Jetz et al., 2012) using the Hackett backbone, implemented via the *consensus* function in the *ape* R package v5.8 (Paradis & Schliep, 2019).

### Land-use metrics

We use forest size as a proxy of forest baseline quality to examine whether small forests are depleted of forest diversity due to species-area effects and human disturbance. We created forest shapefiles and calculated forest size using QGIS v3.22.8 (QGIS Development Team, 2024) based on images provided by Google Earth Pro (U.S. Geological Survey, 2023). Forest shapes were then edited using the Global Forest Change v1.10 (2000-2022) dataset (Hansen et al., 2013) in Google Earth Engine (Gorelick et al., 2017) to obtain forest shapes for the final year of surveying in each study area (e.g. 2018 for Gola). We cross-validated these shapes and sizes with Important Bird & Biodiversity Areas outlined by BirdLife International (2024). When forests consisted of multiple nearby fragments, such as Gola (*n* = 3) and Dompim (*n* = 3), we summed fragment areas to obtain total forest size. While fragmentation can affect species composition, this approach captures total habitat availability at a regional scale, which likely has a greater influence on functional habitat use for many species – particularly in the context of metapopulation dynamics – than fragmentation itself (Bosco et al., 2023; Chasar et al., 2014).

To estimate land-use type and intensity at each survey point, we calculated two continuous quantitative land-use metrics: mean canopy cover percentage and distance to forest edge. Canopy cover is an indicator of local land-use intensity, whereas distance to edge is a measure of proximity to another land-use. We used the Hansen Global Forest Change v1.10 (2000-2022) dataset (Hansen et al., 2013) in Google Earth Engine (Gorelick et al., 2017) to calculate mean canopy cover percentage (henceforth canopy cover) at the time of surveying (i.e. year of survey). To classify individual pixels as forested, we applied a 60% canopy cover threshold, which is considered most accurate at estimating forest loss in tropical forests (Estoque et al., 2021). We calculated canopy cover in a 200 m radius surrounding each survey point which approximately corresponds to the dispersal distance of many rainforest birds (Powell et al., 2015). Using forest shapefiles obtained as described above, we calculated the Euclidean distance of each survey point to the nearest forest (henceforth distance to edge) using the R package *geosphere* v1.5-14 (Hijmans et al., 2021). Distances were adjusted for direction with positive values indicating survey points outside the forest and negative values survey points within the forest. Both metrics were calculated at each survey point (*n* = 2,479) and averaged across survey points pooled to obtain assemblage surveys (*n* = 761).

Using distance to edge to differentiate between forest and agriculture, we removed bird flyovers when these pertained to high overflights of large bird species in inappropriate habitat (for details see Supplementary methods). In each of the two major land-use types (i.e. forest and agriculture), we also omitted species with fewer than three observations in that habitat, as they are likely to contribute minimally to ecosystem functioning and, therefore, are “functionally extinct” in that land-use type (Bregman et al., 2016).

### Quantifying diversity

We calculated species richness (SR), relative abundance (RA), and functional diversity (FD) for six response groups in each assemblage: invertivores, frugivores, granivores, forest birds, forest birds > 500 g, and ant followers. SR was calculated as the total number of species recorded per assemblage. Given that detectability varies between land-use types, particularly between open and closed habitats, we also used RA as a complementary diversity metric. SR is sensitive to the presence of rare or cryptic species, which are more likely to be under-detected in forested sites, potentially leading to an underestimation of biodiversity loss when comparing forests to agricultural landscapes. RA was therefore calculated as the proportion of individuals in a response group relative to the total number of birds recorded in an assemblage survey, controlling for habitat differences in detectability. This approach allowed us to quantify the relative contribution of different functional groups to overall assemblage structure while reducing the influence of uneven detection probabilities across habitats.

While SR and RA provide important measures of taxonomic diversity and community structure, they do not capture the functional differences among species within assemblages. Assemblages with similar species richness may differ greatly in the functional roles represented, particularly if species traits vary in response to land-use change. To assess these differences, we quantified functional diversity (FD), which describes the morphological trait space (“morphospace”) occupied by an assemblage. Specifically, we calculated functional dispersion (FDis), defined as the weighted average distance of individual species to the assemblage centroid in a “morphospace”, as a measure of FD because it is less influenced by outliers than convex-hull measurements of FD, such as functional richness (Laliberte & Legendre, 2010). A higher FDis indicates greater functional dissimilarity among species within an assemblage, potentially also reflecting lower functional redundancy.

Avian functional traits are often highly correlated, primarily due to their association with body size (Pigot et al., 2020). To account for trait collinearity, we followed previous studies (Bregman et al., 2016; Trisos et al., 2014) and applied a two-step principal component analysis (PCA), reducing morphological trait data into three axes linked to ecological functions. Separate PCAs were performed on log-transformed locomotory and trophic traits (pertaining to beak shape), with the first components in both cases correlating strongly with body size. These were combined in a second PCA to produce a single size-related axis, while the second components, representing variation independent of body size, were used as locomotory and trophic trait axes, respectively. We used the three components to calculate FDis in the *fundiversity* package v1.1.1 in R (function *fd_fdis*) (Grenié & Gruson, 2023).

### Data analysis

To assess the effect of forest size on biodiversity, we fitted generalised linear mixed-effects models (GLMMs) for each diversity metric. Forest size was treated as a continuous predictor with SR, RA, and FDis as response variables. For this analysis, we only used data collected within forests. We created models separately for three response groups related to dietary guild (invertivores, frugivores, granivores) and three response groups related to ecological specialisation (forest birds, forest birds > 500 g, ant followers).

We ran a second set of 16 GLMMs to assess the effect of land-use, measured as either canopy cover (scaled and centred) or distance to edge, on SR and FDis. We added an interaction term for forest to measure the difference in biodiversity decline/gain slopes between study areas and carried out post hoc pairwise comparisons between forests using the *emmeans* v1.11.0 R package (Lenth et al., 2024; Searle et al., 1980). To avoid collinearity, we fit models separately for the two measures of land-use and for a subset of four response groups: invertivores, frugivores, forest birds, and forest birds > 500 g. Due to the lack of low-canopy agricultural sampling, we excluded the Gola study from these analyses (see Figure S1-2).

After detecting overdispersion in models fit with a Poisson distribution, all SR models were fit with a generalised Poisson distribution and all RA models were fit with an ordered beta distribution. We used a Gaussian distribution for FDis except for granivores, forest birds > 500 g and ant followers for which we used a Tweedie distribution to address zero-inflation (Lecomte et al., 2013). In all models, we added locality (i.e. landscape within each study area containing multiple spatially clustered transects) as a random effect to account for spatial autocorrelation. To improve model fit, we added season as a predictor rather than random effect to account for temporal autocorrelation (for details see Supplementary methods). All models were fit using the *glmmTMB* v1.1.9 R package (Brooks et al., 2017). We carried out all statistical analyses in R v4.2.2 (R Core Team, 2018).

## Results

### Forest size affects bird assemblage diversity and composition

Invertivore species richness (SR) and relative abundance (RA) did not differ significantly between large and small forests (Figure 3a; Figure S4a; Table S1). However, invertivore functional dispersion (FDis) increased in response to decreasing forest size (Figure S6a). Frugivores diversity declined with forest size consistently across all three metrics (Figure 3b; Figure S4b; Figure S6b), with the largest reduction observed in FDis (̂β: = 0.132, *p* < 0.0001). In contrast, SR, RA and FDis of granivores increased with decreasing forest size (Figure 3c; Figure S4c; Figure S6c; Table S1).

**FIGURE 3.**
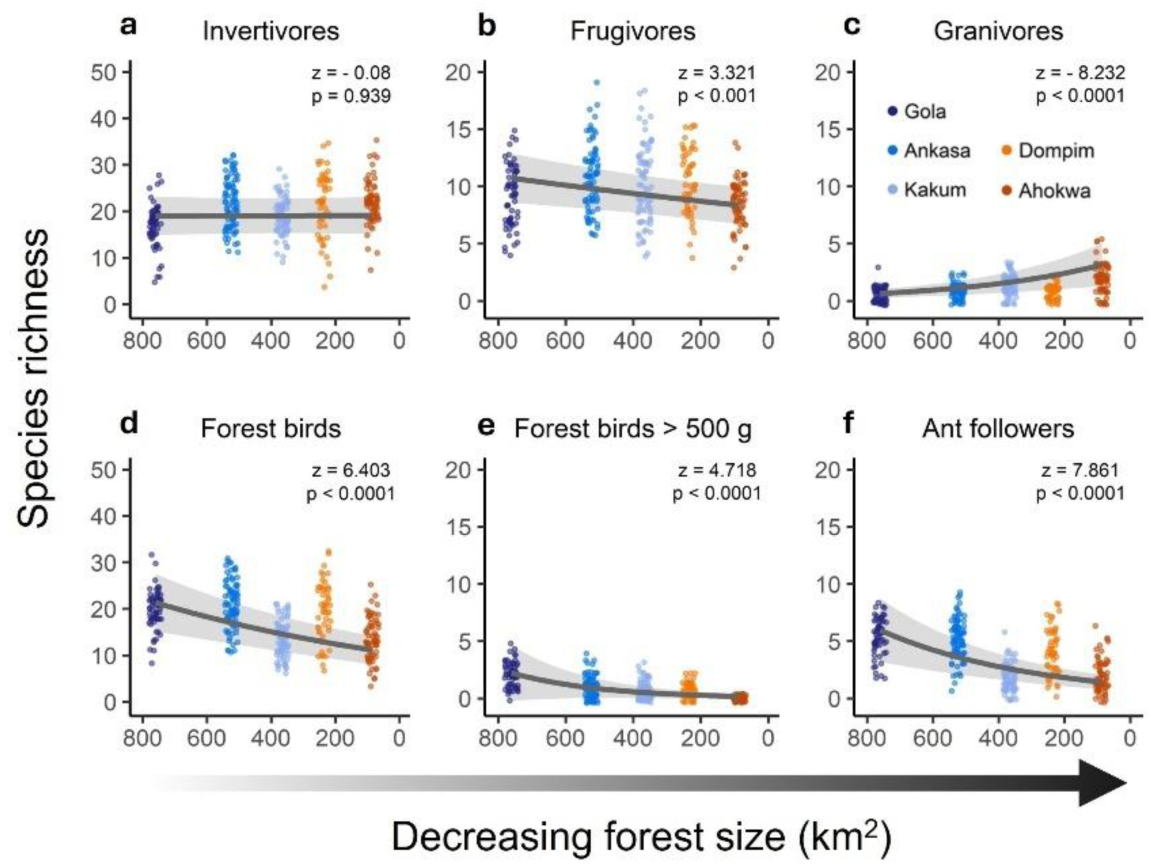
Effect of forest size on the diversity and structure of bird assemblages. Panels (**a-f**) show species richness (coloured dots) detected in surveys of forest assemblages (Gola: *n* = 62; Ankasa: *n* = 83; Kakum: *n* = 75; Dompim: *n* = 53; Ahokwa: *n* = 64). Data are jittered to show distribution of dots. Lines show predictions from univariate mixed effects models assessing the effect of forest size on species richness of forest assemblages. We define forest birds (**d**) as species with high forest dependency and ant followers (**f**) as a subset of specialised forest birds (see Methods). Shaded bands are 95% confidence intervals. Note that y-axis scales vary.

Forest birds showed the steepest decline in RA across response groups (̂β: = 0.489, *p* < 0.0001), constituting an average of 67% of individuals in forest assemblages in the largest forest (Gola), compared to 28% in the smallest forest (Ahokwa; Figure S4d; Table S1). Despite declines in SR (Figure 3d) and RA (Figure S4d), forest bird FDis increased with decreasing forest size (Figure S6d; Table S1), mirroring the pattern observed in invertivore FDis. Ant followers, along with large forest birds, showed significant declines across all three metrics. Notably, large forest birds were recorded at 97% of the forest sites in Gola but were absent from all sites in Ahokwa (0%).

While trends in assemblage-level SR suggested lower richness in smaller forests, total forest SR did not reflect this pattern. Invertivore SR was lowest in Gola, and frugivore and forest bird SR showed little variation across forests (Figure S5). Although species accumulation curves indicated sufficient sampling effort (Figure S1a), additional sightings collected from published reports and citizen-science reveal that up to 139 invertivore species and 58 forest species (73% and 51% of the total invertivore and forest bird community, respectively) may have gone undetected in Gola, making it the forest with the highest total SR (Figure S5). It is difficult to explain this shortfall in sampling completeness, although it may relate to differences in observer experience (Ghana surveys were conducted by professional bird-tour guides with in-depth knowledge of the local avifauna, including songs and calls).

### Forest size mediates bird assemblage responses to land-use change

To investigate whether variations in baseline forest diversity influence gradients of biodiversity loss, we compared the steepness of these gradients from forest to agriculture across four of our study areas (Ankasa, Kakum, Dompim, Ahokwa), using two land-use measures: canopy cover and distance to edge. Invertivore and forest bird SR declined significantly across all forests in response to both decreasing canopy cover and increasing distance from the forest (Figure 4a,c,e,g; Table S2-S3). In contrast, their FDis increased significantly in relation to both predictors in Ankasa, Dompim, and Ahokwa (Figure S7-8; Table S4-S5).

**FIGURE 4.**
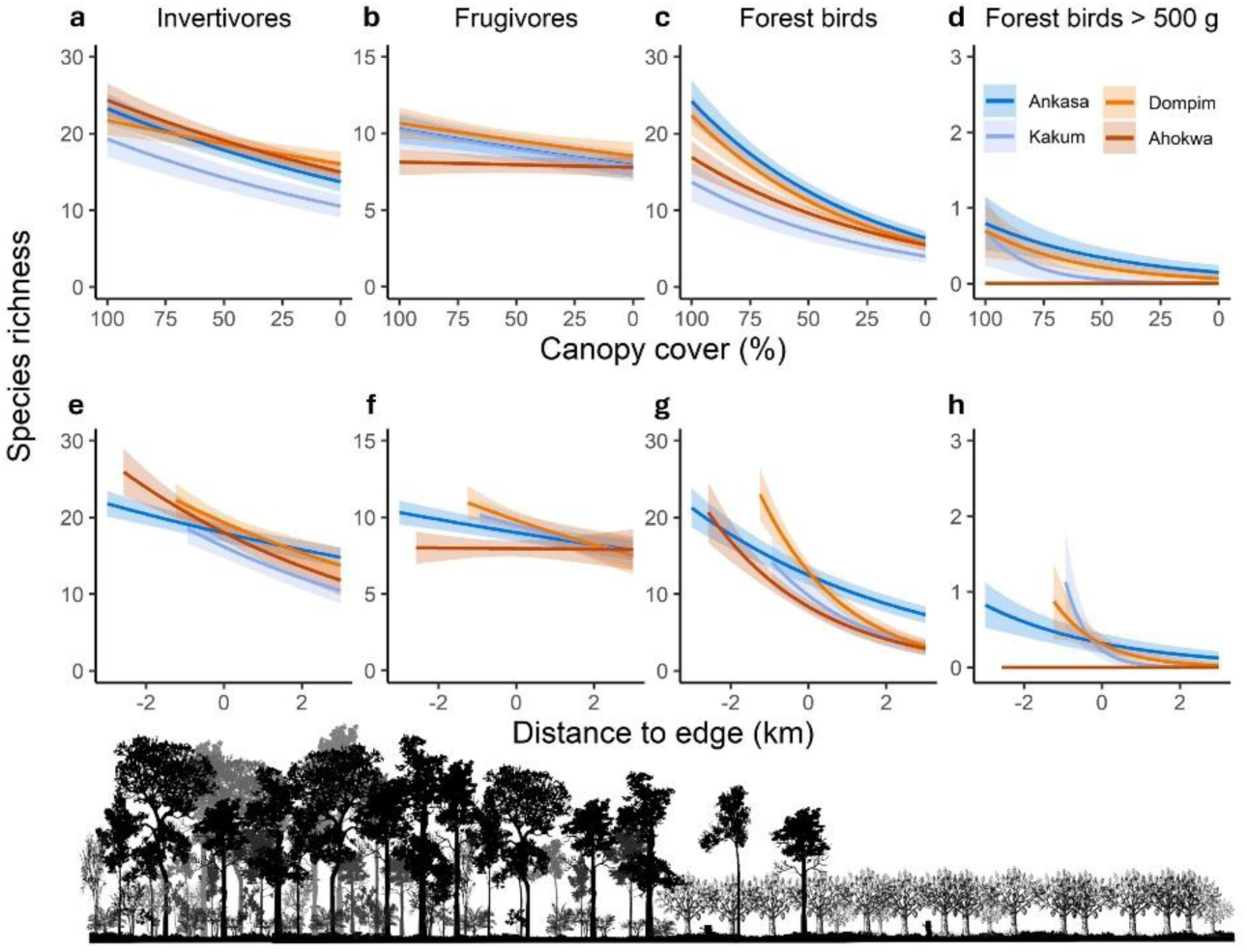
Impact of forest baseline and land-use change on the species richness of bird assemblages. Results are outputs of mixed effect models examining assemblage-level species richness responses to two measures of land-use change: (**a-d**) canopy cover and (**e-h**) distance to forest edge. In **e-h**, zero indicates the forest edge, with negative values indicating distances within the forest and positive values indicating distances outside the forest. Distances to forest edge were capped at 3 km for ease of visualisation. Models were fit with bird assemblage surveys as response variable (Ankasa: *n* = 133; Kakum: *n* = 150; Dompim: *n* = 118; Ahokwa: *n* = 117). Gola was removed from analysis because the landscape surrounding the forest is dominated by agroforestry and therefore not comparable to the landscape context surrounding the other forests (see Methods). Lines indicate minimal adequate model predictions for each study area; shaded bands are 95% confidence intervals. For ease of visualisation error bars of the insignificant slope for Ahokwa (red) were omitted in **d** and **h** and reported in SI (see Table S2-S3). Note that y-axis scales vary.

Frugivore SR declined in the agriculture surrounding the three larger forests (Ankasa, Kakum, Dompim) but showed no change in Ahokwa, where frugivore diversity in forest was already similar to that in agricultural areas (Figure 4b,f; Figure 5b,f; Table S2-S3). Although frugivore SR in agricultural areas was comparable across the four study areas (Figure 4b,f), frugivore assemblages in sites surrounding Ankasa (the largest forest) exhibited higher FDis than those in the agriculture surrounding the other forests (Figure S7f), leading to a shallower decline slope in Ankasa than in the medium-sized forests (Kakum and Dompim) (Figure S8f; Table S5).

**FIGURE 5.**
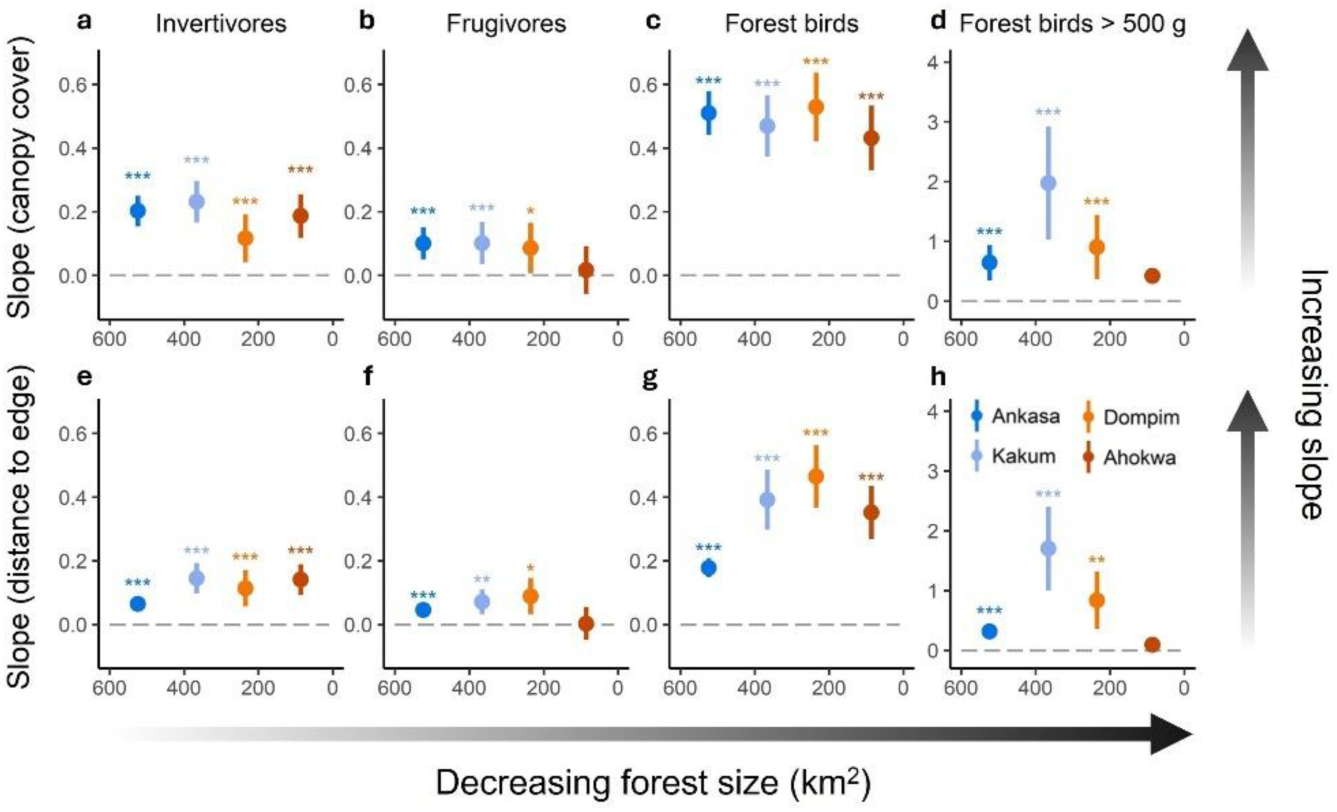
Forest size affects bird assemblage responses to land-use change. Results are outputs of mixed effect models identifying how assemblage-level species richness changes in response to two continuous measures of land-use change: canopy cover (**a-d**) and distance to forest edge (**e-h**). Results shown are coefficient estimates and 95% confidence intervals. Models were fit with bird assemblage surveys as response variable (Ankasa: *n* = 133; Kakum: *n* = 150; Dompim: *n* = 118; Ahokwa: *n* = 117). Gola was removed from analysis because the landscape surrounding the forest is dominated by mixed agroforestry and therefore not comparable to the landscape context surrounding the other forests (see Methods). Slopes are absolute coefficient estimates to facilitate the comparison of steepness (i.e. absolute distance from zero) between slopes. Two contrasting patterns can be observed: large forests have steeper declines of forest-dependent species when canopy cover decreases (**b**,**c**), and small forests have steeper declines in forest-dependent species when the distance to the forest increases (**e**,**g**,**h**). For ease of visualisation, error bars of the non-significant slope for Ahokwa (red) were omitted in **d** and **h** and reported in SI (see Table S2-S3). * = p < 0.05, ** = p < 0.001, *** = p < 0.0001. Note that y-axis scales vary.

While the slopes of decline for invertivores showed little variation between forests with respect to canopy cover (Figure 5a; Table S6), they differed significantly among Ankasa, Kakum, and Ahokwa in response to distance from the edge. Ankasa exhibited a shallower slope, with a higher invertivore SR supported farther from the forest (Figure 5e; Table S6). This pattern was more pronounced for forest birds (Figure 5g), with Ankasa having a significantly shallower slope of decline than all other forests – for example, Dompim (interaction term ̂β: = -0.2875, *p* < 0.0001) (see Table S8 for all pairwise comparisons) – because SR of forest birds at Ankasa remains relatively high in matrix habitats beyond the forest boundary (Figure 4g). Similarly, large forest bird SR declined with decreasing canopy cover (Figure 4d; Table S2) and increasing distance from the forest edge (Figure 4h; Table S3), with steeper declines observed in medium-sized forests Kakum (interaction term ̂β: = -1.386, *p* = 0.0001) and Dompim (interaction term ̂β: = -5.199, *p* = 0.033) compared to Ankasa in response to distance from the edge (Figure 5d,h; Table S9). In Ahokwa, the smallest forest, large forest birds were absent from both agricultural and forest sites, whereas they were recorded in 15% of points at least 1 km from the forest in Ankasa, 8% in Dompim, and 0% in Kakum.

## Discussion

Comparing bird survey data across five West African forests, we found wide variation in assemblage diversity and composition in relation to forest size. Moreover, we show that these shifts in forest baseline assemblages can confound estimates of biodiversity loss in response to land-use change. Specifically, our analyses suggest that biodiversity loss may be systematically underestimated when field studies select either very large and intact baselines, or small and degraded baselines, highlighting the critical importance of baseline selection in space-for-time comparisons.

The main implications of these findings relate to the widespread use of space-for-time comparisons to predict biotic responses to both land-use change (e.g. Cordier et al., 2021; Ferreira et al., 2023; Gray et al., 2016; Millard et al., 2021; Purvis et al., 2018) and climate change (Lovell et al., 2023; Uhler et al., 2021). Comparing across space is often the only method available to assess these responses, yet the results may be biased by a range of factors, including species response lags (Daskalova et al., 2020; Kharouba & Williams, 2024) and changes to local biota over time (Damgaard, 2019; Stouffer et al., 2021), as well as inappropriate baselines (Alstad et al., 2016; Didham et al., 2020; Duarte et al., 2009; Soga & Gaston, 2018). Our study explicitly compares the slope of biodiversity loss across land-use gradients, showing that more-intact baselines provide improved accuracy, except when they are so intact that the sampling gradient may be too short.

Although our study is limited by a relatively small sample of study areas, the patterns we detect suggest that smaller forests have a lower diversity of forest-dependent birds, whereas larger forests have a substantial biodiversity ‘shadow’ resulting from the occurrence of forest species further beyond forest boundaries into the surrounding landscape, extending over the typical length of land-use gradients studied in tropical landscapes. Both of these patterns – i.e. reduced biodiversity in small forests and spillover from species-rich larger forests – can result in shallower slopes of biodiversity decline when comparing natural vegetation to adjacent agriculture. Clearly, any study designed to interpret biotic change by sampling across gradients or other space-for-time comparisons needs to consider the suitability of baselines in the context of landscape history, or to supplement sampling where possible with appropriate time-series data (Dornelas et al., 2018).

### Mechanisms driving shifting baselines

We show that space-for-time comparisons may be biased when small forests are used as the reference baseline since much of their biodiversity (and bioabundance) has already been lost. The negative effects of decreasing forest size on forest-dependent species – large-bodied ones in particular – have been widely documented (Beier et al., 2002; Bovo et al., 2018; Bregman et al., 2014; Sekercioglu et al., 2002). The relatively stable climates of the tropics have driven a range of physiological adaptations to microhabitats which make it harder for tropical forest birds to cope with modified environments. For example, species occurring in the shady understorey of tropical forests are characterised by large eyes, light sensitivity and preference for moist, cool microclimates (Jirinec et al., 2022; Stratford & Robinson, 2005). The interior of small forests tends to be relatively dry, warm and bright compared with larger forests (Laurance et al., 2011), reducing their suitability for many specialised forest species (Stouffer et al., 2021; Stratford & Robinson, 2005). Understory species, in particular, are adversely affected by denser understory vegetation and increased light levels commonly found in small forests (Arcilla et al., 2015; Stouffer et al., 2021). Another factor is low dispersal ability, which reduces population connectivity in fragmented habitats, driving local extinctions in many tropical forest invertivores (Salisbury et al., 2012; Weeks et al., 2023). Meanwhile, most granivores benefit from these altered conditions (Figure 3c), likely because of their greater dispersal ability, coupled with increased habitat permeability and food resources (Donoso et al., 2004).

While changes in landscape configuration and microhabitat are thought to play a prominent role in the decline of small-bodied forest birds (Figure 3d), particularly invertivores (Jirinec et al., 2022; Sekercioglu et al., 2002; Weeks et al., 2023), the decline of large forest birds (Figure 3e) may be caused by a different set of threats. These include patch-size constraints on species with low density populations and large home-range sizes, meaning that smaller habitat patches are not able to support viable population sizes. In addition, larger-bodied species may drop out of smaller forests because of insufficient food resources and increased hunting pressure (Thiollay, 2007; Whytock et al., 2016). For example, the decline we observed in frugivores (Figure 3b) reflects a pattern reported in smaller forest fragments worldwide (Bregman et al., 2014), and which in West Africa is largely driven by the ongoing decline of hornbills and other large frugivorous species, often caused by intensive hunting (Holbech et al., 2018).

Most small-bodied forest birds have small territories or home ranges (Powell et al., 2015), so their populations are less likely to face spatial constraints in smaller forest patches. Ant followers, however, are a notable exception. Ant followers often attend swarms from multiple army ant colonies, which are highly mobile (Stouffer & Bierregaard, 1995). Their home ranges are therefore up to ten times larger than those of other similarly sized forest birds (Martínez et al., 2021; Willson, 2004). Additionally, army ants themselves rely on cool forest interiors, often avoiding hotter and drier conditions characteristic of forest edges and smaller forests (Peters et al., 2008; Peters & Okalo, 2009). Considering the combined effects of microhabitat conditions, home range requirements, and the presence of army ants, it is unsurprising that we observed a particularly steep decline in ant followers within small forests (Figure 3f).

### Biodiversity ‘shadows’: large forests require longer sampling gradients

It may seem counter-intuitive to expect a shallower decline of biodiversity when the baseline starts from a higher point. However, biases in space-for-time comparisons may also emerge when selecting large forests as baselines because of their very high diversity and different landscape history (Figure 4e,g,h). This pattern could potentially arise from at least two different mechanisms. First, the most extensive forests support larger source populations, more effectively buffering against biodiversity loss in surrounding landscapes through source-sink dynamics, whereby a higher diversity of forest-dependent birds disperse beyond the boundaries of large forests into agricultural areas. Second, larger forests are associated with more remote regions, less populated by humans, and therefore with a shorter timeline and lower intensity of habitat change in the surrounding landscape. These factors may all lead to remnant populations of forest birds surviving, at least temporarily, in partially cleared landscapes surrounding the largest forests and extending further from the forest boundary. Consequently, biodiversity in agriculture or agroforestry near large forests may be boosted by source-sink dynamics and landscape history. Our results suggest that survey transects should be lengthened to account for this effect, in the case of very intact baselines, so that transects extend far enough into intensive land-uses and beyond the forest’s biodiversity ‘shadow.’

### Establishing ecological impacts of land-use change in the context of shifting baselines

We find that some ecological and functional groups are far more sensitive to land-use change than others, consistent with the results of previous studies in tropical forest systems (Arcilla et al., 2015; Bregman et al., 2016; Jarrett et al., 2021; Stouffer et al., 2021). Although responses differed between groups, each group showed a consistent response pattern to both canopy cover and distance to edge, with invertivores, frugivores and forest birds – large forest birds in particular – declining in response to land-use change. While the direction of response was consistent within each group, the steepness of the slope varied between the two land-use metrics. This was especially evident in forest birds, where declines in response to distance to edge were shallower in larger forests, but declines in response to canopy cover remained steep regardless of forest size (Figure 5c,g). Our findings are consistent with previous research suggesting that agricultural areas with sufficient canopy cover and proximity to extensive natural forest can maintain populations of forest birds (Ferreira et al., 2023; Jarrett et al., 2021; Olimpi et al., 2022), including species providing important services, such as insect predation (Bouarakia et al., 2023; Olimpi et al., 2022; Yahya et al., 2024) and seed dispersal (Bregman et al., 2016).

The critical importance of extensive forest blocks is further supported by the steep declines we detect in the functional dispersion of frugivores as forest size decreases, suggesting that species with key traits, such as wide beaks required for dispersing large-seeded trees (Galetti et al., 2013; Michalski et al., 2007), are likely to be lost in smaller forests and their surrounding agriculture (Bovo et al., 2018; Caves et al., 2013). A particular example highlighted by our study, is the absence of large hornbills in the smallest forest we surveyed, suggesting severe erosion of seed dispersal services for large-seeded trees (de Assis Bomfim et al., 2018; Holbech et al., 2018).

Focusing on forest birds in general, and invertivores in particular, we found the opposite pattern of functional dispersion remaining stable or increasing in smaller, more fragmented forests, as noted in some previous studies (Coster et al., 2015; Luck et al., 2013; Magnago et al., 2014). This suggests that disturbed forests can support functionally diverse communities, often because they are colonised by non-forest species with divergent traits, in the case of invertivores. However, when the increase in functional dispersion is paired with a decline in species richness, as we found for forest birds, this indicates a loss of functionally similar species and, consequently, a decrease in functional redundancy. Although declining numbers of species with similar roles may not immediately affect ecosystem functioning, lower levels of functional redundancy may reduce ecosystem resilience because there are fewer surplus species to provide an ‘insurance effect’ in the context of changing environmental conditions and further species loss (Fonseca & Ganade, 2001; Naeem, 1998). Our findings highlight the role of large, intact forests in supporting ecosystem functions and services in adjacent agricultural areas, but also the need for more nuanced biodiversity assessments that consider landscape history, source-sink dynamics and functional resilience when evaluating land-use impacts.

### Caveats and limitations

Our analyses focus on changes in biodiversity occurring in recent decades, yet baselines also shift over much longer timeframes. Even if we could sample biodiversity in our study areas centuries ago, animal communities worldwide were already degraded at that point due to Pleistocene extinctions shifting what is perceived as “natural” in modern times (Søndergaard et al., 2025). With the caveat that we are focusing on relatively recent baselines, we have shown substantial differences in avian diversity at the level of local assemblages depending on forest size. Nonetheless, we note that overall species richness was relatively constant across forest blocks (Figure S5), suggesting that most forest species survive even in small forests. Our results highlight that many such species only survive in tiny or patchy remnant populations, contributing little to ecological processes given their local rarity, and facing high risk of local extinction.

Whether species that went undetected in our surveys are either very rare or locally extinct is difficult to determine given that some tropical forest birds are hard to detect because they are silent or unobtrusive (Robinson et al., 2018). The apparent disappearance of many large forest-associated species in smaller habitat patches is likely to reflect local extinctions, although small populations of wary – and therefore less detectable – individuals may survive. Either way, further sampling or better detection rates in our surveys would almost certainly increase the number of large bird species identified in the most intact forest blocks, further amplifying the differences observed between small and large forests, thereby strengthening our main conclusions. Ultimately, the collapse of some forest bird populations over recent decades has removed their contributions to ecological processes, whether they survive in small numbers or not (Bregman et al., 2016; Bregman et al. 2014).

Differences in the slopes of biodiversity loss across our study areas may also reflect climatic and environmental factors, including variation in rainfall. For instance, the largest Ghanaian forest block we studied (Ankasa) lies in the southwest of Western Region, an area characterised by higher rainfall, while the smallest site (Ahokwa) is located in the drier northeast of Western Region. Drier conditions may exacerbate area and edge effects, reducing populations of core forest species in habitat patches. In addition, patterns of biodiversity loss may be influenced by forest shape, with narrow and more-fragmented forest blocks losing biodiversity more rapidly than circular habitat patches. The vegetation of the surrounding matrix can also be influential, with certain types of agricultural land-uses supporting more forest species than others (e.g. Hatfield et al., 2020; Sanderson et al., 2022). Finally, the historical timeline of forest loss plays a role, with levels of biodiversity remaining high in recently deforested landscapes where extinction debts have yet to be paid in full (Daskalova et al., 2020). Future studies should account for these factors where possible to provide a more nuanced understanding of biodiversity loss after land-use change and the accuracy of space-for-time comparisons.

## Conclusion

Our study shows that biodiversity trends derived from space-for-time comparisons are influenced by the chosen reference baseline. A degraded baseline may lead to underestimates of biodiversity loss because many species have already declined or disappeared in the face of altered microhabitat conditions, human exploitation, and demographic constraints on species with low density populations. Similarly, biodiversity loss may even be underestimated from highly intact baselines associated with unpaid extinction debts and more extensive overspill of biodiversity into surrounding landscapes, suggesting that sampling gradients should be lengthened in this context. Although this study focuses on birds, the underlying mechanisms have wider relevance across many animal and plant systems, highlighting the potential risks of space-for-time comparisons more generally. Future studies should incorporate landscape history, regional and historical biodiversity data, and other contextual information to enable more accurate estimates of baseline biodiversity prior to recent environmental change.

## Acknowledgements

We thank Williams K. Apraku, Beth Downe, Mike Ford, Robert Ntakor, Robert O. Appau, Ben Phalan, Felicity Edwards, Mohamed S. Konneh, Patrick Dauda, Robin Whytock, Andy Schofield, Denis Bannah, Aruna Senesie, Alfried P. Vogler and Luke L. Powell for help with fieldwork, data or useful discussions. This research was funded by the UKRI Global Challenges Research Fund grant ES/P011306/1 (JAT). Fieldwork in Kakum was funded by the Natural Environment Research Council grants NE/P001092/1 and NE/P00394X/1, and the Ecosystem Services for Poverty Alleviation (ESPA) Program grant NE/K010379-1. Fieldwork in Gola was funded by the Darwin Initiative grant 20-022. AD was supported by a Natural Environment Research Council studentship through the Science and Solutions for a Changing Planet Doctoral Training Partnership.

## SUPPLEMENTARY INFORMATION

### Supplementary methods

#### Study region and selection of study areas

The Upper Guinean rainforest once extended across West Africa from Guinea and Sierra Leone in the west through Liberia, Côte d’Ivoire and Ghana to Togo in the east, where it is separated from the Lower Guinean rainforest by a tract of drier savannah habitat (the Dahomey Gap). Vast areas of the original forest have been cleared and fragmented, leaving only remnant patches of natural habitat, mostly within protected areas, which continue to be threatened by agricultural expansion, logging and mining (Mittermeier et al., 1999). To assess impacts of land-use change on fragments of Upper Guinean rainforest, we focused our study on the Western Region of Ghana. This region has a tropical monsoonal climate with a mean annual precipitation > 1500 mm and mean annual temperature of 26°C (Atiah et al., 2020).

We carried out bird surveys along forest-to-agriculture gradients in three study areas (Ankasa, Dompim, Ahokwa) selected to represent different forest sizes and levels of intactness (Figure 2a). Ankasa Game Reserve is a large (524 km^2^) protected forest, while Dompim (235 km^2^) is a recently fragmented forest that still retains relatively intact, tall-stature patches. Ahokwa, the smallest forest block (86 km^2^), has been fragmented for over 20 years and is undergoing extensive logging and conversion to agriculture along its edges (Figure S3).

To increase sample size, we then conducted a systematic literature survey and selection process to identify further studies with compatible design across West Africa. We initially compiled a dataset of seven case studies comparing avian diversity between Afrotropical rainforest baselines and their surrounding agriculture. After excluding five of the studies because of sampling differences (see Data S1), we included two additional protected sites – Gola and Kakum – with compatible methods and habitat types to our surveys. Gola Rainforest National Park in Sierra Leone (761 km²) represents one of the largest remaining tracts of intact forest in the region, with minimal recent disturbance and well-preserved tall-stature canopy cover (Klop et al., 2010). Kakum National Park in Ghana (366 km²) underwent extensive selective logging until 1989, resulting in a lower canopy height compared to the other protected areas included in our study (Ankasa, Gola, see Figure S3). However, following a logging ban in 1989, Kakum has supported increasingly regenerating and structurally complex closed canopy forest again (Wiafe, 2016).

#### Survey methods

In Ankasa, Dompim and Ahokwa, we used a two-tier stratified random sampling design to select study areas within and adjacent to each forest block. Initially, each study area was overlaid with a spatially referenced 1 km × 1 km grid, dividing the area into potential study cells. We then categorised each grid cell into one of four broad land classifications based on the dominant land type, using forest cover data from Hansen et al. (2013) via Google Earth Engine (Gorelick et al., 2017). These classifications included forest interior (> 95% forest cover), forest edge (25–75% forest cover), agriculture (< 5% forest cover), and other (e.g. urban areas or large water bodies). For the purpose of this study, only cells classified as forest interior, forest edge, or agriculture were included in the sampling frame.

From each of these land classes, 30% of grid cells were randomly selected for bird sampling. During fieldwork, some of these sites were found to be inaccessible, requiring further modifications to the sampling design to account for logistical constraints and the need to obtain permissions from local communities. In cases where selected cells were found to be inaccessible, adjustments were made to the sampling plan on the day. Within the final selection of grid cells, we established survey transects along existing paths or roads, extending these where necessary by cutting trails into the forest. Where straight-line transects were not feasible due to vegetation, waterways, or terrain, we used curved or angled transects, with the curvature and degree of angle minimised.

Along each survey transect, we selected survey points in the relevant habitat, ensuring a minimum spacing of 200 m between points. Bird surveys were conducted at each point, using a standardised point-count method. Surveys started at dawn to align with peak bird activity. On arrival at each point, observers remained quiet for one minute to reduce any disturbance caused by their presence, then identified all birds seen or heard during a 10-minute count. Observers estimated the number of individuals encountered for each species, minimising the double counting of individuals or groups. Flyovers (i.e. birds not using the vegetation within a 50-m radius from the survey point) were recorded separately. Once a count was completed, observers moved to the next survey point and repeated the process. In total, we used these methods to survey bird communities at 1,222 point-count sites along 116 transects (Ankasa: *n* = 42; Dompim: *n* = 39; Ahokwa: *n* = 35).

In these three study areas, we removed flyovers of large-bodied species not using the relevant habitat at the survey point – particularly where these species were detected in habitats unlikely to support them, such as waterbirds flying over agricultural fields. This decision was based on the likelihood that, in highly fragmented landscapes, wide-ranging species may traverse unsuitable habitats between habitat fragments, resulting in detections that reflect movement through the landscape rather than actual habitat use. Overall, we removed 19 total observations, comprising 12 flyover records of three large forest species (*Bycanistes fistulator*, *Bycanistes subcylindricus*, *Pteronetta hartlaubii*) from agricultural surveys, and 7 observations of two large non-forest species (*Corvus albus*, *Milvus aegyptius*) from forest surveys. It was not necessary to apply this adjustment to Gola or Kakum data because flyovers were excluded from the survey protocol.

Both additional study areas – Gola and Kakum – used survey methods broadly similar to those described above. In Gola, bird surveys were conducted over six field seasons (four dry and two wet) between 2013 and 2018. Surveys took place along 35 transects, each comprising 7 to 26 point-count sites spaced at a minimum of 200 m. Transects extended up to 4 km into Gola Rainforest National Park and up to 27 km into the surrounding agricultural landscape. Point counts were conducted between 06:45 and 11:15. During each count, all birds seen or heard were recorded, with observers estimating cluster size for each species. Surveys in Gola were longer than those at our other sites, with a duration of 15 minutes compared to 10 minutes elsewhere. However, even in tropical forests, the majority of individuals – typically over 90% – are detected within the first 5 to 10 minutes of a count (Lee & Marsden, 2008; Mattos & Peris, 2008). Based on this, we treated Gola surveys as comparable to the others when grouping adjacent point-count sites into assemblage surveys. A total of 817 point-count sites were surveyed across 35 transects.

In Kakum, bird surveys were conducted over five field seasons (three wet and two dry) between 2014 and 2017. Surveys were carried out along three forest-to-agriculture gradients, beginning in Kakum National Park and extending into the surrounding cocoa farming communities of Aboabo, Homaho, and Kwameameobang. Along each gradient, transects consisting of 3 points each, were established at regular distances from the forest edge. The innermost transect was located 1 km inside the forest, while the outermost extended 5 km into the agricultural landscape. Survey points along each transect were spaced at least 200 m apart. Each point was surveyed once per field season using standardised 10-minute point counts, with most surveys taking place between 6:15 and 11:30 and a small set of surveys carried out at dusk (*n* = 69). During each count, observers recorded all birds seen or heard, noting species identity and estimated cluster size. In total, 440 point-count sites were surveyed across 150 transects.

#### Resolving taxonomic differences between datasets

Differences in avian taxonomy across datasets can pose a challenge when integrating records from multiple studies. The three most commonly used taxonomies – BirdLife International (2024), eBird based on the Clements checklist (Clements et al., 2023), and BirdTree (Jetz et al., 2012) – do not always align, with some splitting a single species into multiple geographically distinct daughter lineages. Both the Gola and Kakum datasets used the BirdTree taxonomy (Jetz et al., 2012) which enabled us to follow an established taxonomy crosswalk (Tobias et al., 2022) to resolve discrepancies and standardise all species-level records across studies. We added corresponding BirdLife and eBird taxonomy names to all records. Where a species was split differently across taxonomies, we used eBird geographical range maps to examine the ranges of all daughter lineages and selected the one overlapping with our study sites. For instance, the Chestnut Wattle-eye (*Platysteira castanea*) from the BirdTree taxonomy is divided into two lineages according to BirdLife/eBird taxonomies: the West African Wattle-eye (*Dyaphorophyia* [*Platysteira*] *hormophora*), ranging from Ghana to Guinea and the Chestnut Wattle-eye (*Dyaphorophyia* [*Platysteira*] *castanea*), found from southern Benin to northern Zambia. In this case, we matched all *Platysteira castanea* records to *Dyaphorophyia hormophora* (BirdLife) and *Platysteira hormophora* (eBird), as it is the only daughter lineage with a geographical range overlapping our study sites. This approach allowed us to integrate all data into a consistent taxonomy, resulting in a unified dataset of 46,481 bird records across all sites.

### Forest dependency

We extracted forest dependency data from BirdLife International (2024) for all species in our dataset (*n* = 413), classifying them as high (*n* = 58), medium (*n* = 196) or low (*n* = 99) forest dependency and as non-forest (*n* = 60) if not normally occurring in forest. However, these forest dependency data are based on a relatively crude system that mainly relies on researchers with little field experience of the species or areas involved estimating forest dependency from text in Birds of the World (Billerman et al., 2022). This approach introduces substantial uncertainty and may underestimate forest dependency for two main reasons.

First, literature sources used to compile the summary information in species accounts of Birds of the World (Billerman et al., 2022) may mention occurrence of forest species in clearings, farmland, gardens or other non-forest habitats based on relatively few published observations, or records from well-visited sites adjacent to extensive forests. These infrequent observations are then often summarised into a list of habitats from which a species is known, which can be misinterpreted as evidence of general habitat use by species with high dependence on forest. For example, species such as Fraser’s Eagle Owl *Bubo poensis* (= *Ketupa poensis*) and Great Blue Turaco *Corythaeola cristata* are listed by BirdLife International as medium forest dependency whereas their populations are largely restricted to forest and its immediate vicinity.

Second, descriptions of habitat use in Birds of the World (Billerman et al., 2022) are drawn from information across the global range of species, which introduces further uncertainty because some bird species vary geographically in their habitat niche, or occupy a wider range of habitats in some regions than others. For example, the Western Crested Guineafowl (*Guttera verreauxi*) is classified as medium forest dependency by BirdLife International on the basis of occurrence in non-forest habitat in parts of its geographical distribution, whereas the West African form (*G. v. verreauxi*) is largely restricted to forest.

To correct for these biases, we reviewed BirdLife International forest dependency classifications using literature and eBird, an online repository of locality data. Where recent observations of species are predominantly clustered around major forest patches in the Upper Guinean rainforest, we assumed that species had a high level of forest dependency, at least in West Africa. Following this procedure, we changed the forest dependency of 65 highly forest-associated species, from medium to high forest dependency (for specific cases, see Data S1).

### Ant followers

The focus on ant followers stems from their ecological specialisation, which makes them highly sensitive to habitat disturbance and spatial constraints (Martínez et al., 2021; Peters et al., 2008; Stouffer & Bierregaard, 1995). Unlike other small-bodied forest birds with relatively small home ranges (Powell et al., 2015), ant followers typically have larger territories, often up to ten times the size of similarly sized forest species (Martínez et al., 2021; Willson, 2004). These birds attend multiple mobile colonies of army ants, which rely on cool forest interiors and avoid the hotter, drier conditions at forest edges (Peters & Okalo, 2009). Consequently, ant followers are especially vulnerable to declines in smaller forests, making them an important group for investigating the effects of forest size and structure on forest birds.

While the distinction between obligate and facultative ant followers is well documented in the Neotropics (e.g. Martínez et al., 2021; Swartz, 2001; Willson, 2004), this classification remains poorly studied in the Afrotropics (Craig, 2022). To date, only three Afrotropical species – *Alethe castanea* (closely related to *Alethe diademata* in our dataset), *Chamaetylas poliocephala*, and *Neocossyphus poensis* – have been identified as likely obligate ant followers, based on behaviours such as bivouac checking, which is typically associated with obligate ant followers in the Neotropics (Nikolaou et al., 2025; Rodrigues, 2024). Due to the limited availability of detailed behavioural data, we classified all species that habitually attend army ant swarms as ant followers. We used Billerman et al. (2022) as an initial source to identify species within our dataset frequently reported to attend ant swarms. Species reported as ant followers without reference to frequency of attendance were noted as potential ant followers. We excluded species reported only occasionally at swarms – such as *Ispidina picta* and *Ispidina lecontei* (Billerman et al., 2022) – despite their inclusion as ant followers in other studies (Craig, 2022).

We refined these classifications by cross-referencing additional published literature (e.g. Peters et al., 2008; Waltert et al., 2024; Willis, 1983) and adjusting our categorisations accordingly. We reclassified *Eurillas latirostris* as a non-ant follower due to inconsistent evidence regarding its frequency at swarms; some sources describe it as a frequent attendant (Peters et al., 2008), while others report only occasional attendance (Waltert et al., 2024). In contrast, we classified *Picathartes gymnocephalus* as an ant follower based on consistent reports of regular attendance (Thompson & Fotso, 2000; Willis, 1983). We also classified *Criniger barbatus* as an ant follower, based on evidence of regular ant-following by a closely related sister species, *Criniger chloronotus* (Billerman et al., 2022; Waltert et al., 2024).

### Canopy height

Canopy cover and distance to edge are standard metrics for assessing land use and proximity to natural habitats. However, they do not capture levels of disturbance within forested habitat. In the remaining fragments of Upper Guinean forest, managed and illegal logging occurs at varying intensities, typically targeting the tallest and oldest trees (Hansen et al., 2012; Hansen & Treue, 2008). To estimate forest condition, we therefore calculated mean canopy height in a 200 m radius surrounding each survey point (henceforth canopy height) using Google Earth Engine (Gorelick et al., 2017). Canopy height layers for our study areas were obtained from the GLAD Global Forest Canopy Height dataset (Potapov et al., 2022).

### Accounting for temporal autocorrelation

We conducted two sets of analyses: the first to assess the impact of forest size on bird diversity, and the second to evaluate the influence of land use (measured by canopy cover or distance to edge) on bird diversity. Model fit for each analysis was assessed using the *DHARMa* R package (Hartig, 2022). To account for temporal autocorrelation, we included sampling season as a fixed effect in all models. Season was treated as a fixed effect, rather than a random effect, to address residual heteroskedasticity. For models exploring the effect of forest size on the functional dispersion of granivores and large forest birds, season was excluded to resolve model convergence issues.

In the first set of analyses, examining forest size, season as a fixed effect accounted for temporal autocorrelation both between the dry and wet seasons, and across years to capture long-term population trends. In contrast, in the second set of analyses examining the effects of land-use change, we used a fixed effect for season (dry/wet) but did not include year of sampling, to improve model fit.

## Supplementary tables

**Table S1.** Effect of forest size on species richness, relative abundance and functional dispersion of bird assemblages. Results shown are model summaries predicting the effect of forest size on bird response groups. For details on model structure see Methods and Supplementary methods. * = p < 0.05, ** = p < 0.001, *** = p < 0.0001.

|  | Response group | Estimate | Standard error | z-value | p-value |  |
| --- | --- | --- | --- | --- | --- | --- |
| **Species richness** | Invertivores | -0.0019 | 0.0249 | -0.08 | 0.9398 |  |
|  | Frugivores | 0.0838 | 0.0252 | 3.321 | < 0.001 | ** |
|  | Granivores | -0.5305 | 0.0644 | -8.232 | < 0.0001 | *** |
|  | Forest birds | 0.2177 | 0.034 | 6.403 | < 0.0001 | *** |
|  | Forest birds > 500 g | 0.8378 | 0.1776 | 4.718 | < 0.0001 | *** |
|  | Ant followers | 0.4771 | 0.0607 | 7.861 | < 0.0001 | *** |
| **Relative abundance** | Invertivores | 0.0251 | 0.0478 | 0.526 | 0.599 |  |
|  | Frugivores | 0.194 | 0.0447 | 4.339 | < 0.0001 | *** |
|  | Granivores | -0.5669 | 0.0876 | -6.472 | < 0.0001 | *** |
|  | Forest birds | 0.4887 | 0.0565 | 8.653 | < 0.0001 | *** |
|  | Forest birds > 500 g | 1.1373 | 0.258 | 4.399 | < 0.0001 | *** |
|  | Ant followers | 0.5578 | 0.0741 | 7.531 | < 0.0001 | *** |
| **Functional dispersion** | Invertivores | -0.108 | 0.0192 | -5.633 | < 0.0001 | *** |
|  | Frugivores | 0.1318 | 0.0249 | 5.293 | < 0.0001 | *** |
|  | Granivores | -1.0068 | 0.1657 | -6.078 | < 0.0001 | *** |
|  | Forest birds | -0.0884 | 0.025 | -3.534 | < 0.001 | ** |
|  | Forest birds > 500 g | 1.4763 | 0.3274 | 4.509 | < 0.0001 | *** |
|  | Ant followers | 0.5202 | 0.0748 | 6.797 | < 0.0001 | *** |

**Table S2.** Impact of canopy cover on the species richness of bird assemblages. Results shown are model summaries predicting the effect of canopy cover on the species richness of bird response groups. For details on model structure see Methods and Supplementary methods. * = p < 0.05, ** = p < 0.001, *** = p < 0.0001.

| Predictor | Response group | Case study | Estimate | Std error | z-value | p-value |  |
| --- | --- | --- | --- | --- | --- | --- | --- |
| Canopy cover | Invertivores | Ankasa | 0.2022 | 0.0249 | 8.1329 | < 0.0001 | *** |
|  |  | Kakum | 0.2314 | 0.0223 | 10.3942 | < 0.0001 | *** |
|  |  | Dompim | 0.1165 | 0.0288 | 4.0395 | < 0.0001 | *** |
|  |  | Ahokwa | 0.1865 | 0.0245 | 7.6242 | < 0.0001 | *** |
|  | Frugivores | Ankasa | 0.1004 | 0.0259 | 3.8808 | < 0.0001 | *** |
|  |  | Kakum | 0.1011 | 0.0218 | 4.6296 | < 0.0001 | *** |
|  |  | Dompim | 0.0857 | 0.0309 | 2.7757 | < 0.01 | * |
|  |  | Ahokwa | 0.0168 | 0.028 | 0.6009 | 0.5479 |  |
|  | Forest birds | Ankasa | 0.5097 | 0.0352 | 14.4858 | < 0.0001 | *** |
|  |  | Kakum | 0.4693 | 0.0343 | 13.6687 | < 0.0001 | *** |
|  |  | Dompim | 0.5287 | 0.0418 | 12.6568 | < 0.0001 | *** |
|  |  | Ahokwa | 0.4314 | 0.0381 | 11.3092 | < 0.0001 | *** |
|  | Forest birds |  |  |  |  |  |  |
|  | > 500 g | Ankasa | 0.6449 | 0.151 | 4.2698 | < 0.0001 | *** |
|  |  | Kakum | 1.9764 | 0.4583 | 4.3125 | < 0.0001 | *** |
|  |  | Dompim | 0.9039 | 0.2303 | 3.9245 | < 0.0001 | *** |
|  |  | Ahokwa | 0.4062 | 6575.3 | < 0.0001 | 0.9999 |  |

**Table S3.** Impact of distance to forest edge on the species richness of bird assemblages. Results shown are model summaries predicting the effect of distance to edge on the species richness of bird response groups. For details on model structure see Methods and Supplementary methods. * = p < 0.05, ** = p < 0.001, *** = p < 0.0001.

| Predictor | Response group | Case study | Estimate | Std error | z-value | p-value |  |
| --- | --- | --- | --- | --- | --- | --- | --- |
| **Distance to edge** | **Invertivores** | Ankasa | -0.0645 | 0.0085 | -7.5253 | < 0.0001 | *** |
|  |  | Kakum | -0.1452 | 0.0226 | -6.4128 | < 0.0001 | *** |
|  |  | Dompim | -0.114 | 0.0278 | -4.0953 | < 0.0001 | *** |
|  |  | Ahokwa | -0.141 | 0.023 | -6.1416 | < 0.0001 | *** |
|  | **Frugivores** | Ankasa | -0.0455 | 0.0082 | -5.4929 | < 0.0001 | *** |
|  |  | Kakum | -0.071 | 0.0185 | -3.8242 | < 0.001 | ** |
|  |  | Dompim | -0.0889 | 0.0281 | -3.1589 | < 0.01 | * |
|  |  | Ahokwa | -0.0026 | 0.0248 | -0.1066 | 0.9151 |  |
|  | **Forest birds** | Ankasa | -0.1777 | 0.0149 | -11.9272 | < 0.0001 | *** |
|  |  | Kakum | -0.3918 | 0.0457 | -8.5738 | < 0.0001 | *** |
|  |  | Dompim | -0.4652 | 0.0473 | -9.8359 | < 0.0001 | *** |
|  |  | Ahokwa | -0.3519 | 0.0396 | -8.8821 | < 0.0001 | *** |
|  | **Forest birds** |  |  |  |  |  |  |
|  | **> 500 g** | Ankasa | -0.3157 | 0.0628 | -5.0244 | < 0.0001 | *** |
|  |  | Kakum | -1.7015 | 0.3509 | -4.8494 | < 0.0001 | *** |
|  |  | Dompim | -0.8356 | 0.2356 | -3.5472 | < 0.001 | ** |
|  |  | Ahokwa | 0.0085 | 4757 | < 0.0001 | 0.9999 |  |

**Table S4.** Impact of canopy cover on the functional dispersion of bird assemblages. Results shown are model summaries predicting the effect of canopy cover on the functional dispersion of bird response groups. For details on model structure see Methods and Supplementary methods. * = p < 0.05, ** = p < 0.001, *** = p < 0.0001.

| Predictor | Response group | Case study | Estimate | Std error | z-value | p-value |  |
| --- | --- | --- | --- | --- | --- | --- | --- |
| **Canopy cover** | **Invertivores** | Ankasa | -0.1558 | 0.0187 | -8.3142 | < 0.0001 | *** |
|  |  | Kakum | 0.0179 | 0.0144 | 1.2402 | 0.2149 |  |
|  |  | Dompim | -0.108 | 0.0221 | -4.8737 | < 0.0001 | *** |
|  |  | Ahokwa | -0.0631 | 0.0191 | -3.3067 | < 0.001 | ** |
|  | **Frugivores** | Ankasa | 0.1393 | 0.0194 | 7.1849 | < 0.0001 | *** |
|  |  | Kakum | 0.1123 | 0.016 | 7.0288 | < 0.0001 | *** |
|  |  | Dompim | 0.1563 | 0.0233 | 6.7057 | < 0.0001 | *** |
|  |  | Ahokwa | 0.032 | 0.02 | 1.5975 | 0.1101 |  |
|  | **Forest birds** | Ankasa | -0.1451 | 0.0297 | -4.8843 | < 0.0001 | *** |
|  |  | Kakum | -0.0096 | 0.0261 | -0.3689 | 0.7122 |  |
|  |  | Dompim | -0.0762 | 0.0364 | -2.0908 | < 0.05 | * |
|  |  | Ahokwa | -0.133 | 0.0311 | -4.2787 | < 0.0001 | *** |
|  | **Forest birds** |  |  |  |  |  |  |
|  | **> 500 g** | Ankasa | 0.7913 | 0.366 | 2.1617 | < 0.05 | * |
|  |  | Kakum | 2.4066 | 1.1245 | 2.1402 | < 0.05 | * |
|  |  | Dompim | 2.1712 | 1.4832 | 1.4638 | 0.1432 |  |
|  |  | Ahokwa | -2.839 | 50733 | < 0.0001 | 0.9999 |  |

**Table S5.**
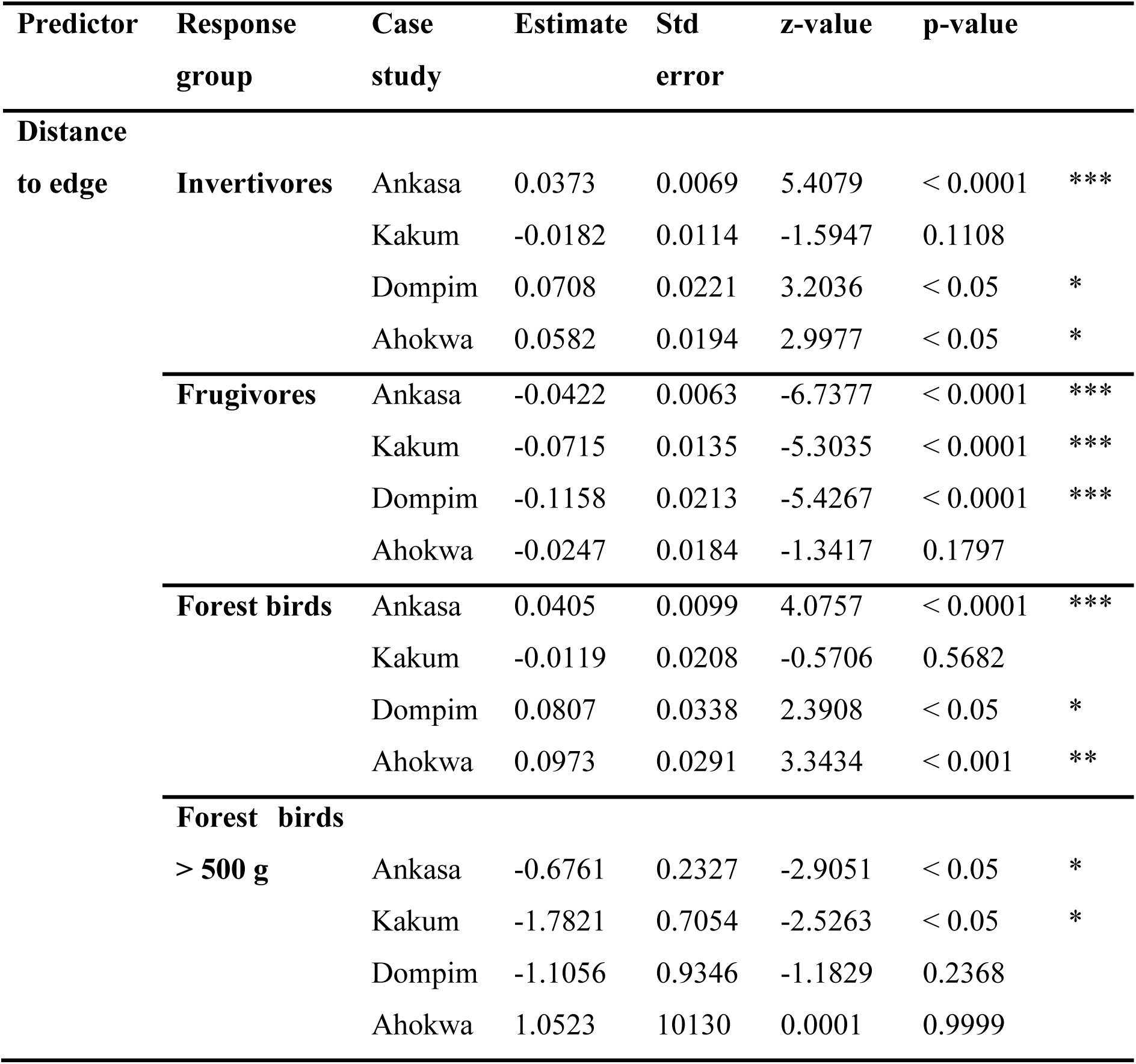
Impact of distance to forest edge on the functional dispersion of bird assemblages. Results shown are model summaries predicting the effect of distance to edge on the functional dispersion of bird response groups. For details on model structure see Methods and Supplementary methods. * = p < 0.05, ** = p < 0.001, *** = p < 0.0001.

| Predictor | Response group | Case study | Estimate | Std error | z-value | p-value |  |
| --- | --- | --- | --- | --- | --- | --- | --- |
| **Distance to edge** | **Invertivores** | Ankasa | 0.0373 | 0.0069 | 5.4079 | < 0.0001 | *** |
|  |  | Kakum | -0.0182 | 0.0114 | -1.5947 | 0.1108 |  |
|  |  | Dompim | 0.0708 | 0.0221 | 3.2036 | < 0.05 | * |
|  |  | Ahokwa | 0.0582 | 0.0194 | 2.9977 | < 0.05 | * |
|  | **Frugivores** | Ankasa | -0.0422 | 0.0063 | -6.7377 | < 0.0001 | *** |
|  |  | Kakum | -0.0715 | 0.0135 | -5.3035 | < 0.0001 | *** |
|  |  | Dompim | -0.1158 | 0.0213 | -5.4267 | < 0.0001 | *** |
|  |  | Ahokwa | -0.0247 | 0.0184 | -1.3417 | 0.1797 |  |
|  | **Forest birds** | Ankasa | 0.0405 | 0.0099 | 4.0757 | < 0.0001 | *** |
|  |  | Kakum | -0.0119 | 0.0208 | -0.5706 | 0.5682 |  |
|  |  | Dompim | 0.0807 | 0.0338 | 2.3908 | < 0.05 | * |
|  |  | Ahokwa | 0.0973 | 0.0291 | 3.3434 | < 0.001 | ** |
|  | **Forest birds** |  |  |  |  |  |  |
|  | **> 500 g** | Ankasa | -0.6761 | 0.2327 | -2.9051 | < 0.05 | * |
|  |  | Kakum | -1.7821 | 0.7054 | -2.5263 | < 0.05 | * |
|  |  | Dompim | -1.1056 | 0.9346 | -1.1829 | 0.2368 |  |
|  |  | Ahokwa | 1.0523 | 10130 | 0.0001 | 0.9999 |  |

**Table S6.** Comparison of invertivore slopes across forest baselines in response to land-use change. Results shown are interaction terms from linear mixed-effects models assessing the impact of land-use change, measured by canopy cover and distance to edge, on invertivore species richness and functional dispersion. For details on the model structure, refer to the Supplementary methods. Slopes that are significantly different between forests are denoted with * = p < 0.05, ** = p < 0.001, *** = p < 0.0001.

| **Invertivores** |  |  |  |  |  |  |
| --- | --- | --- | --- | --- | --- | --- |
| **Diversity metric** | **Predictor** | **Forest comparison** | **Estimate** | **Std error** | **z-value** | **p-value** |
| **Species richness** | **Canopy cover** | Ankasa – Kakum | 0.0291 | 0.033 | 0.8728 | 0.3828 |
|  |  | Ankasa – Dompim | -0.0857 | 0.038 | -2.2413 | 0.025 * |
|  |  | Ankasa – Ahokwa | -0.0157 | 0.035 | -0.4505 | 0.6523 |
|  |  | Kakum – Dompim | -0.1149 | 0.036 | -3.1544 | 0.0016 * |
|  |  | Kakum – Ahokwa | -0.0449 | 0.033 | -1.3559 | 0.1751 |
|  |  | Dompim – Ahokwa | 0.07 | 0.038 | 1.85 | 0.0643 |
|  | **Distance to edge** | Ankasa – Kakum | -0.0808 | 0.024 | -3.3353 | 0.0008 ** |
|  |  | Ankasa – Dompim | -0.0495 | 0.029 | -1.7018 | 0.0888 |
|  |  | Ankasa – Ahokwa | -0.0766 | 0.025 | -3.1237 | 0.0018 * |
|  |  | Kakum – Dompim | 0.0313 | 0.036 | 0.8715 | 0.3835 |
|  |  | Kakum – Ahokwa | 0.0042 | 0.032 | 0.1294 | 0.897 |
|  |  | Dompim – Ahokwa | -0.027 | 0.036 | -0.7509 | 0.4527 |
| **Functional dispersion** | **Canopy cover** | Ankasa – Kakum | 0.1778 | 0.024 | 7.3415 | < 0.0001 *** |
|  |  | Ankasa – Dompim | 0.0479 | 0.029 | 1.6597 | 0.0969 |
|  |  | Ankasa – Ahokwa | 0.0928 | 0.027 | 3.4667 | 0.0005 ** |
|  |  | Kakum – Dompim | -0.1259 | 0.026 | -4.7598 | < 0.0001 *** |
|  |  | Kakum – Ahokwa | -0.081 | 0.024 | -3.3844 | 0.0007 ** |
|  |  | Dompim – Ahokwa | 0.0449 | 0.029 | 1.5342 | 0.125 |
|  | **Distance to edge** | Ankasa – Kakum | -0.0555 | 0.013 | -4.1593 | < 0.0001 *** |
|  |  | Ankasa – Dompim | 0.0335 | 0.023 | 1.4492 | 0.1473 |
|  |  | Ankasa – Ahokwa | 0.0209 | 0.021 | 1.0164 | 0.3094 |
|  |  | Kakum – Dompim | 0.089 | 0.025 | 3.5785 | 0.0003 ** |
|  |  | Kakum – Ahokwa | 0.0764 | 0.023 | 3.3923 | 0.0007 ** |
|  |  | Dompim – Ahokwa | -0.0126 | 0.029 | -0.428 | 0.6687 |

**Table S7.** Comparison of frugivore slopes across forest baselines in response to land-use change. Results shown are interaction terms from linear mixed-effects models assessing the impact of land-use change, measured by canopy cover and distance to edge, on frugivore species richness and functional dispersion. For details on the model structure, refer to the Supplementary methods. Slopes that are significantly different between forests are denoted with * = p < 0.05, ** = p < 0.001, *** = p < 0.0001.

| **Frugivores** |  |  |  |  |  |  |
| --- | --- | --- | --- | --- | --- | --- |
| **Diversity metric** | **Predictor** | **Forest comparison** | **Estimate** | **Std error** | **z-value** | **p-value** |
| **Species richness** | **Canopy cover** | Ankasa – Kakum | 0.0007 | 0.034 | 0.0203 | 0.9839 |
|  |  | Ankasa – Dompim | -0.0146 | 0.04 | -0.3629 | 0.7166 |
|  |  | Ankasa – Ahokwa | -0.0836 | 0.038 | -2.1941 | 0.0282 |
|  |  | Kakum – Dompim | -0.0153 | 0.038 | -0.4046 | 0.6858 |
|  |  | Kakum – Ahokwa | -0.0842 | 0.036 | -2.373 | 0.0177 |
|  |  | Dompim – Ahokwa | -0.069 | 0.042 | -1.6533 | 0.0983 |
|  | **Distance to edge** | Ankasa – Kakum | -0.0254 | 0.02 | -1.2507 | 0.211 |
|  |  | Ankasa – Dompim | -0.0434 | 0.029 | -1.4776 | 0.1395 |
|  |  | Ankasa – Ahokwa | 0.0429 | 0.026 | 1.642 | 0.1005 |
|  |  | Kakum – Dompim | -0.0179 | 0.034 | -0.5316 | 0.595 |
|  |  | Kakum – Ahokwa | 0.0683 | 0.031 | 2.2072 | 0.0273 |
|  |  | Dompim – Ahokwa | 0.0862 | 0.037 | 2.3017 | 0.0214 |
| **Functional dispersion** | **Canopy cover** | Ankasa – Kakum | -0.0271 | 0.025 | -1.0766 | 0.2816 |
|  |  | Ankasa – Dompim | 0.017 | 0.03 | 0.5621 | 0.574 |
|  |  | Ankasa – Ahokwa | -0.1073 | 0.028 | -3.8476 | 0.0001 |
|  |  | Kakum – Dompim | 0.0441 | 0.028 | 1.5587 | 0.1191 |
|  |  | Kakum – Ahokwa | -0.0802 | 0.026 | -3.1295 | 0.0017 |
|  |  | Dompim – Ahokwa | 0.0449 | 0.029 | 1.5342 | 0.125 |
|  | **Distance to edge** | Ankasa – Kakum | -0.0555 | 0.013 | -4.1593 | < 0.0001 |
|  |  | Ankasa – Dompim | 0.0335 | 0.023 | 1.4492 | 0.1473 |
|  |  | Ankasa – Ahokwa | 0.0209 | 0.021 | 1.0164 | 0.3094 |
|  |  | Kakum – Dompim | 0.089 | 0.025 | 3.5785 | 0.0003 |
|  |  | Kakum – Ahokwa | 0.0765 | 0.023 | 3.3923 | 0.0007 |
|  |  | Dompim – Ahokwa | -0.0126 | 0.029 | -0.428 | 0.6687 |

**Table S8.** Comparison of forest bird slopes across forest baselines in response to land-use change. Results shown are interaction terms from linear mixed-effects models assessing the impact of land-use change, measured by canopy cover and distance to edge, on forest bird species richness and functional dispersion. For details on the model structure, refer to the Supplementary methods. Slopes that are significantly different between forests are denoted with * = p < 0.05, ** = p < 0.001, *** = p < 0.0001.

| Forest birds |  |  |  |  |  |  |  |
| --- | --- | --- | --- | --- | --- | --- | --- |
| Diversity metric | Predictor | Forest comparison | Estimate | Std error | z-value | p-value |  |
| Species richness | Canopy cover | Ankasa – Kakum | -0.0404 | 0.049 | -0.8217 | 0.4112 |  |
|  |  | Ankasa – Dompim | 0.019 | 0.055 | 0.3483 | 0.7276 |  |
|  |  | Ankasa – Ahokwa | -0.0783 | 0.052 | -1.5084 | 0.1314 |  |
|  |  | Kakum – Dompim | 0.0594 | 0.054 | 1.0974 | 0.2725 |  |
|  |  | Kakum – Ahokwa | -0.0379 | 0.051 | -0.7384 | 0.4603 |  |
|  |  | Dompim – Ahokwa | -0.0973 | 0.057 | -1.7179 | 0.0858 |  |
|  | Distance to edge | Ankasa – Kakum | -0.2141 | 0.048 | -4.4564 | < 0.0001 | *** |
|  |  | Ankasa – Dompim | -0.2875 | 0.05 | -5.7811 | < 0.0001 | *** |
|  |  | Ankasa – Ahokwa | -0.1742 | 0.042 | -4.1235 | < 0.0001 | *** |
|  |  | Kakum – Dompim | -0.0733 | 0.066 | -1.1149 | 0.2649 |  |
|  |  | Kakum – Ahokwa | 0.0399 | 0.06 | 0.6612 | 0.5085 |  |
|  |  | Dompim – Ahokwa | 0.0863 | 0.037 | 2.3017 | 0.0214 | * |
| Functional dispersion | Canopy cover | Ankasa – Kakum | 0.1355 | 0.04 | 3.4266 | 0.0006 | ** |
|  |  | Ankasa – Dompim | 0.0469 | 0.047 | 1.4716 | 0.1411 |  |
|  |  | Ankasa – Ahokwa | 0.0121 | 0.043 | 0.2818 | 0.7781 |  |
|  |  | Kakum – Dompim | -0.0666 | 0.045 | -1.4852 | 0.1375 |  |
|  |  | Kakum – Ahokwa | -0.1234 | 0.041 | -3.0411 | 0.0024 | * |
|  |  | Dompim – Ahokwa | -0.0568 | 0.048 | -1.187 | 0.2352 |  |
|  | Distance to edge | Ankasa – Kakum | -0.0523 | 0.023 | -2.2704 | 0.0232 | * |
|  |  | Ankasa – Dompim | 0.04027 | 0.035 | 1.1445 | 0.2524 |  |
|  |  | Ankasa – Ahokwa | 0.0568 | 0.307 | 1.8479 | 0.0646 |  |
|  |  | Kakum – Dompim | 0.0926 | 0.04 | 2.3307 | 0.0198 | * |
|  |  | Kakum – Ahokwa | 0.1091 | 0.036 | 3.0512 | 0.0023 | * |
|  |  | Dompim – Ahokwa | 0.0165 | 0.044 | 0.3714 | 0.7104 |  |

**Table S9.** Comparison of large forest bird slopes across forest baselines in response to land-use change. Results shown are interaction terms from linear mixed-effects models assessing the impact of land-use change, measured by canopy cover and distance to edge, on large forest bird species richness and functional dispersion. For details on the model structure, refer to the Supplementary methods. Slopes that are significantly different between forests are denoted with * = p < 0.05, ** = p < 0.001, *** = p < 0.0001.

| Forest birds > 500 g |  |  |  |  |  |  |
| --- | --- | --- | --- | --- | --- | --- |
| Diversity metric | Predictor | Forest comparison | Estimate | Std error | z-value | p-value |
| Species richness | Canopy cover | Ankasa – Kakum | 1.3314 | 0.483 | 2.7573 | 0.0058 * |
|  |  | Ankasa – Dompim | 0.259 | 0.275 | 0.9408 | 0.3468 |
|  |  | Ankasa – Ahokwa | -0.2205 | 5450 | < 0.0001 | 0.9999 |
|  |  | Kakum – Dompim | -1.0725 | 0.513 | -2.0905 | 0.0366 * |
|  |  | Kakum – Ahokwa | -5.9413 | 51175 | < 0.0001 | 0.9999 |
|  |  | Dompim – Ahokwa | -0.7088 | 5274 | < 0.0001 | 0.9999 |
|  | Distance to edge | Ankasa – Kakum | -1.3858 | 0.357 | -3.8853 | 0.0001 ** |
|  |  | Ankasa – Dompim | -5.1994 | 0.244 | -2.1345 | 0.0328 * |
|  |  | Ankasa – Ahokwa | 0.4109 | 4277 | 9.6045 | 0.9999 |
|  |  | Kakum – Dompim | 0.8659 | 0.423 | 2.0475 | 0.04 * |
|  |  | Kakum – Ahokwa | 2.6643 | 5193 | 0.0005 | 0.9999 |
|  |  | Dompim – Ahokwa | 0.1948 | 5370 | 3.6269 | 0.9999 |
| Functional dispersion | Canopy cover | Ankasa – Kakum | 1.6154 | 1.183 | 1.3651 | 0.1722 |
|  |  | Ankasa – Dompim | 1.3799 | 1.529 | 0.9029 | 0.367 |
|  |  | Ankasa – Ahokwa | 1.0694 | 19856 | 5.3855 | 0.9999 |
|  |  | Kakum – Dompim | -0.2354 | 1.86 | -0.1266 | 0.8993 |
|  |  | Kakum – Ahokwa | -5.8578 | 71597 | < 0.0001 | 0.9999 |
|  |  | Dompim – Ahokwa | -8.6575 | 13607 | < 0.0001 | 0.9999 |
|  | Distance to edge | Ankasa – Kakum | -1.106 | 0.743 | -1.4894 | 0.1364 |
|  |  | Ankasa – Dompim | -0.4295 | 0.963 | -0.446 | 0.6556 |
|  |  | Ankasa – Ahokwa | 2.4867 | 12737 | 0.0001 | 0.9998 |
|  |  | Kakum – Dompim | 0.6765 | 1.17 | 0.5784 | 0.563 |
|  |  | Kakum – Ahokwa | 4.2503 | 20596 | 0.0002 | 0.9998 |
|  |  | Dompim – Ahokwa | 6.4537 | 41586 | 0.0001 | 0.9998 |

## Supplementary figures

**FIGURE S1.**
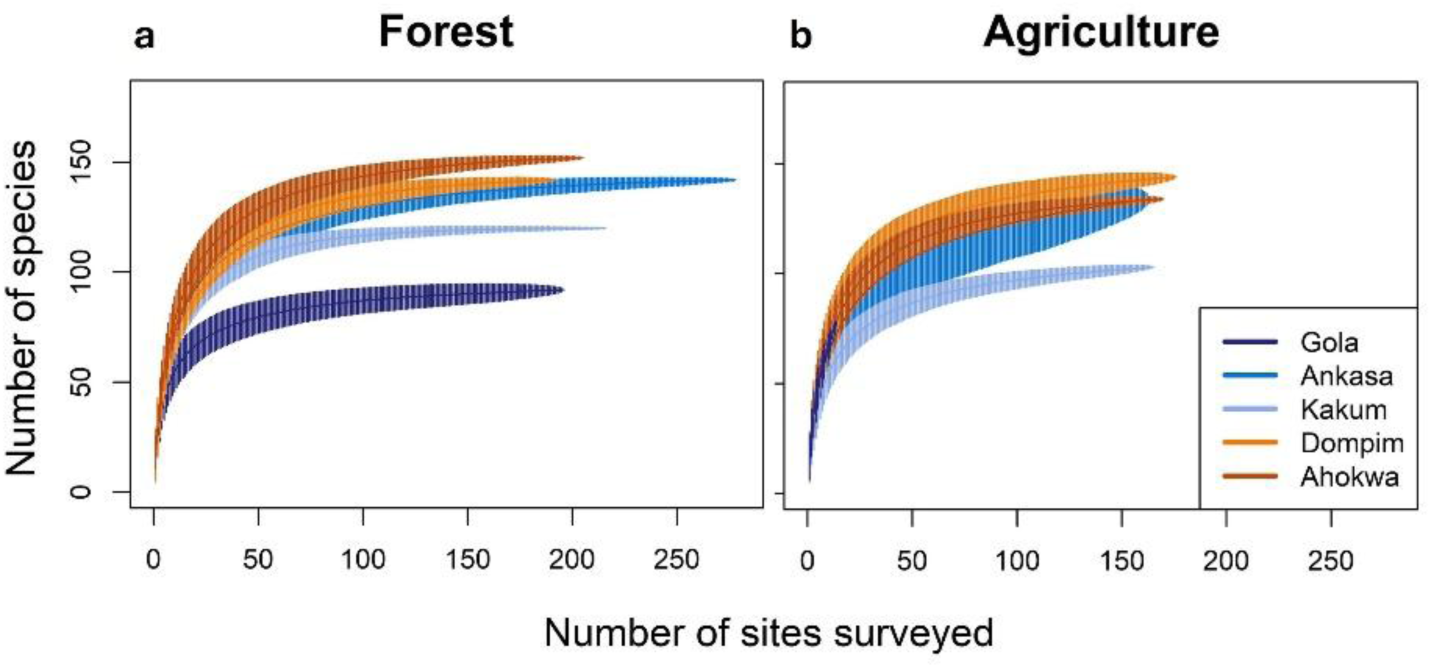
Species accumulation curves show insufficient sampling in agriculture surrounding Gola. Results are species accumulation curves with 95% confidence intervals for sites in (**a**) forest and (**b**) agriculture for each case study (shown in different colours). We define as forest, survey points within forest blocks (as shown in Figure 2a), and as agriculture, survey points outside forest blocks with canopy cover ≤ 40% to exclude high canopy cover agroforestry sites. Study areas can be considered sufficiently sampled if the cumulative species richness ceases to increase when further points are surveyed (i.e. the curve reaches a horizontal asymptote).

**FIGURE S2.**
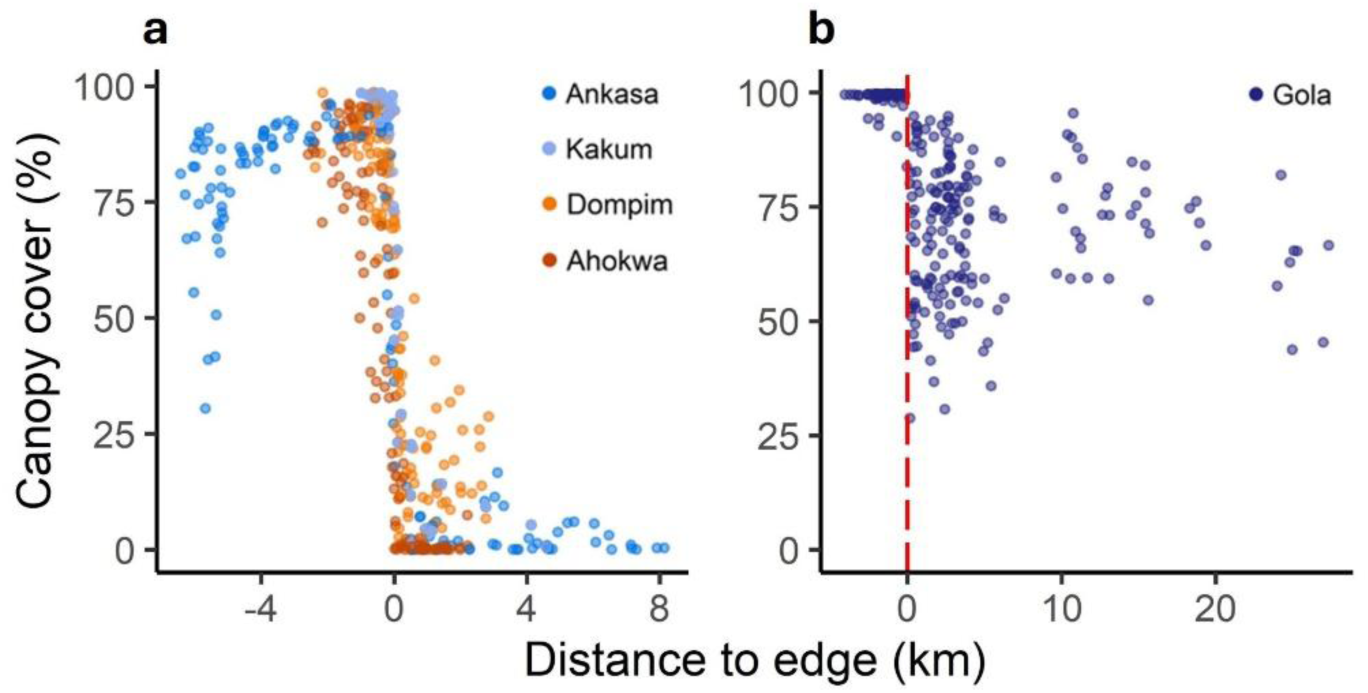
Landscape matrix outside forest varies between case studies. Canopy cover percentage is plotted against distance to forest edge. Dots are coloured by case study and represent assemblage surveys (see Methods). On the x-axis, zero indicates the forest edge, with negative values indicating distances within the forest and positive values indicating distances outside the forest. (**a**) The landscape surrounding Ankasa (*n* = 133), Kakum (*n* = 150), Dompim (*n* = 118), and Ahokwa (*n* = 117) is dominated by low canopy cover agriculture, making canopy cover a useful metric to differentiate between sites within and outside the forest. (**b**) Gola (*n* = 240) is surrounded by agroforestry with high canopy cover. Given its different landscape context, Gola was excluded from analyses comparing forest bird assemblages to agricultural bird assemblages (Figures 4 and 5).

**FIGURE S3.**
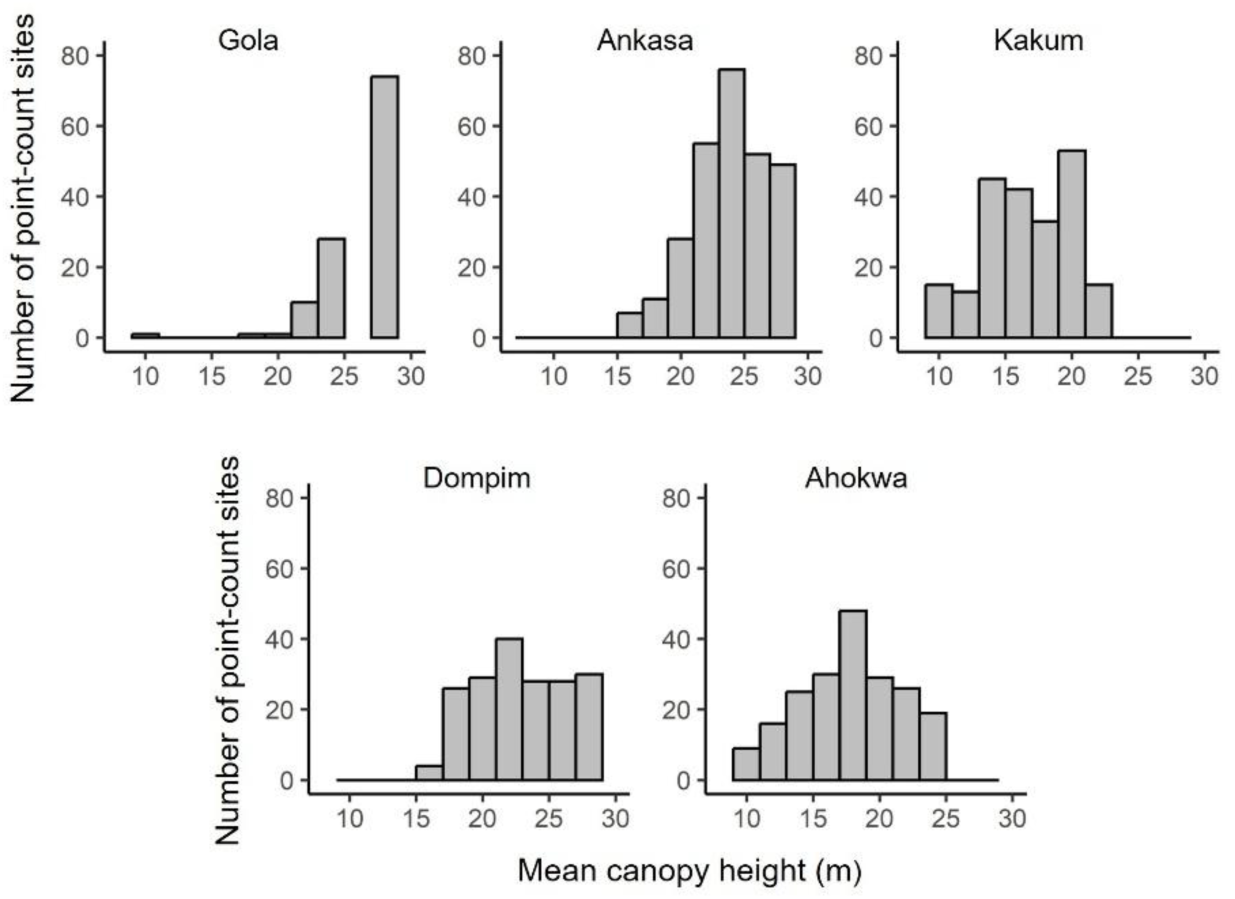
Variation in canopy height across different study areas. Mean canopy height is calculated for all forest point-count sites (Gola: *n* = 196; Ankasa: *n* = 278; Kakum: *n* = 216; Dompim: *n* = 192; Ahokwa: *n* = 205). Kakum and Ahokwa lack point-count sites with canopy height > 25 m.

**FIGURE S4.**
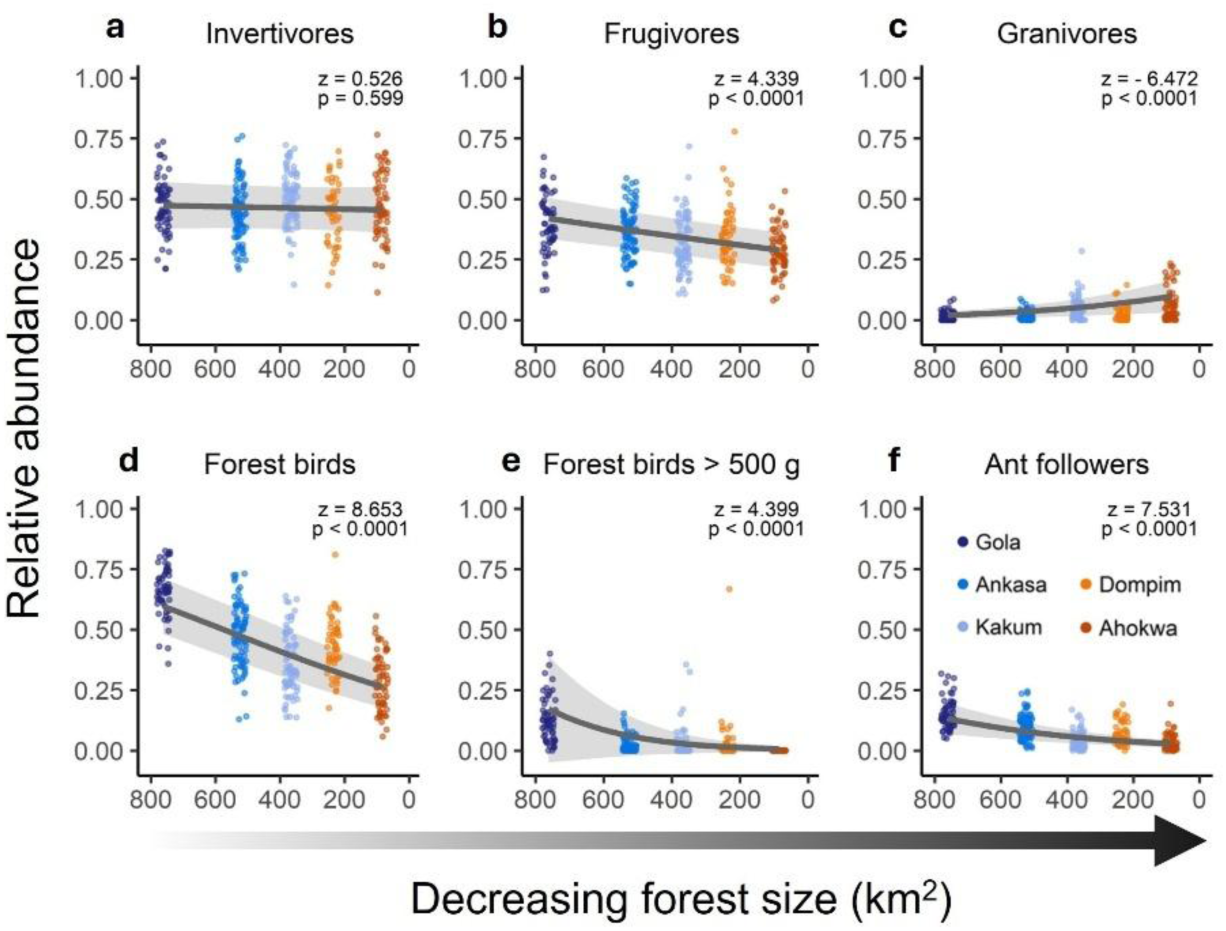
Effect of forest size on the diversity and structure of bird assemblages. Panels (**a-f**) show relative abundance (coloured dots) of different ecological groups in forest assemblages (Gola: *n* = 62; Ankasa: *n* = 83; Kakum: *n* = 75; Dompim: *n* = 53; Ahokwa: *n* = 64). Lines show predictions from univariate mixed effects models assessing the effect of forest size on the relative abundance of specialised bird groups in forest assemblages; shaded bands are 95% confidence intervals. We define forest birds (**d**) as forest-exclusive species and ant followers (**f**) as a subset of specialised forest birds which prey on animals flushed by *Dorylus* army ants (see Methods). Note that y-axis scales vary.

**FIGURE S5.**
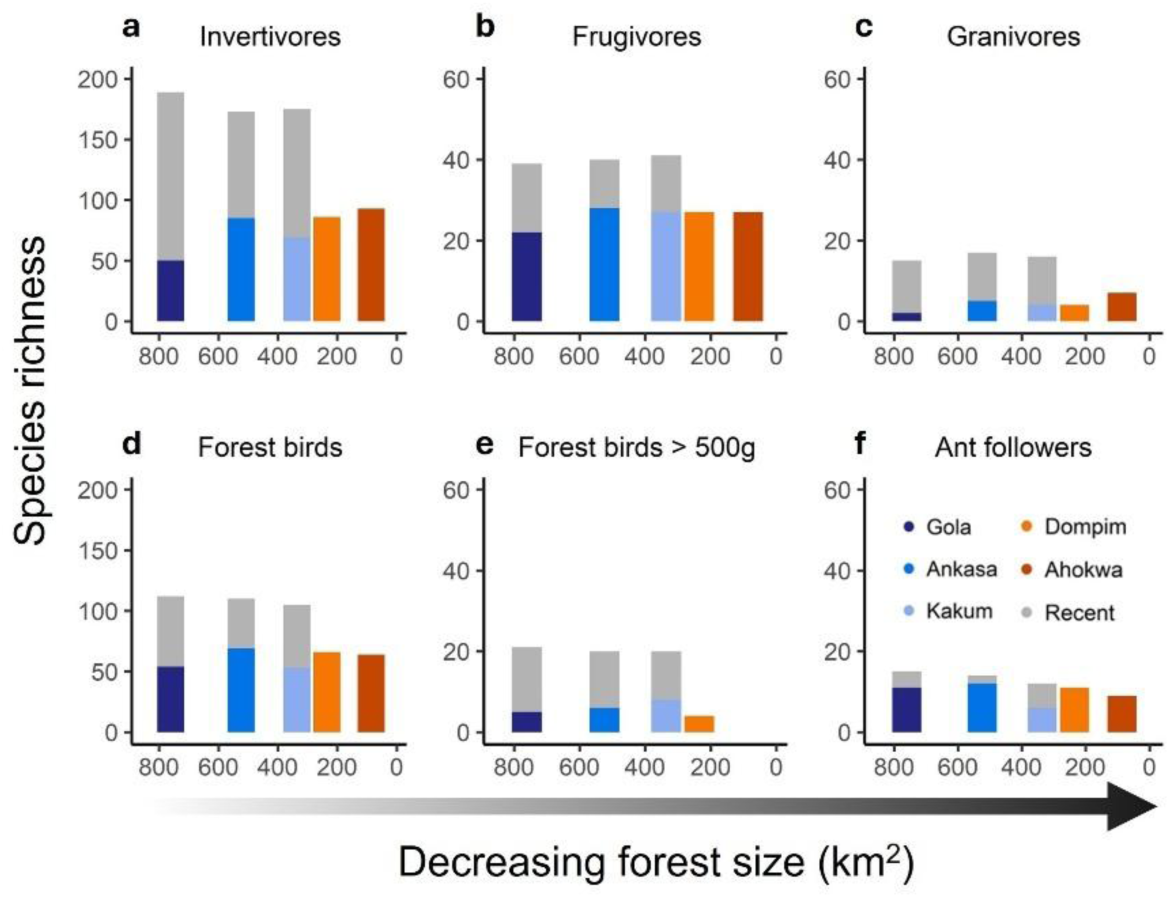
Evaluating completeness of field surveys by comparing with total bird diversity. Panels (**a-f**) show cumulative species richness (coloured bars) for different ecological groups sampled across all point-count sites classified as forest in each study area (Gola: *n* = 62; Ankasa: *n* = 83; Kakum: *n* = 75; Dompim: *n* = 53; Ahokwa: *n* = 64). Grey bars represent additional species richness within the same ecological groups reported by other published field surveys and independent observers (see Methods). These additional data were only available for protected areas (Gola, Ankasa, Kakum). We define forest birds (**d**) as species with high forest dependency and ant followers (**f**) as a subset of specialised forest birds which prey on animals flushed by *Dorylus* army ants (see Methods). Note that y-axis scales vary.

**FIGURE S6.**
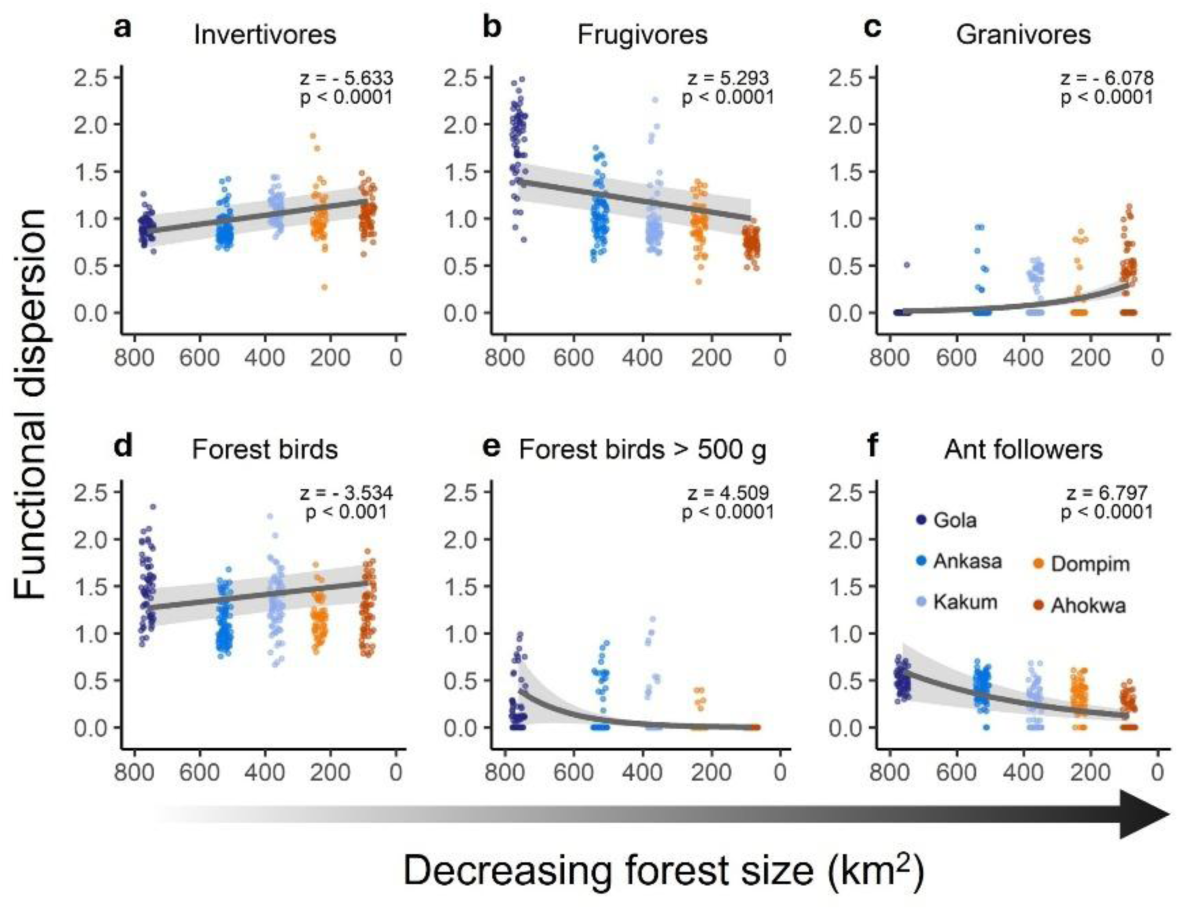
Effect of forest size on the functional diversity of bird assemblages. Plots (**a-f**) show association between forest size and functional dispersion (FDis), defined as the weighted average distance of individual species to the assemblage centroid in a morphospace (Laliberte & Legendre, 2010). Coloured dots represent forest assemblages (Gola: *n* = 62; Ankasa: *n* = 83; Kakum: *n* = 75; Dompim: *n* = 53; Ahokwa: *n* = 64). Lines show predictions from univariate mixed effects models assessing the effect of forest size on FDis of forest assemblages; shaded bands are 95% confidence intervals. We define forest birds (**d**) as forest-exclusive species and ant followers (**f**) as a subset of specialised forest birds which follow *Dorylus* army ants for foraging. Note that y-axis scales vary.

**FIGURE S7.**
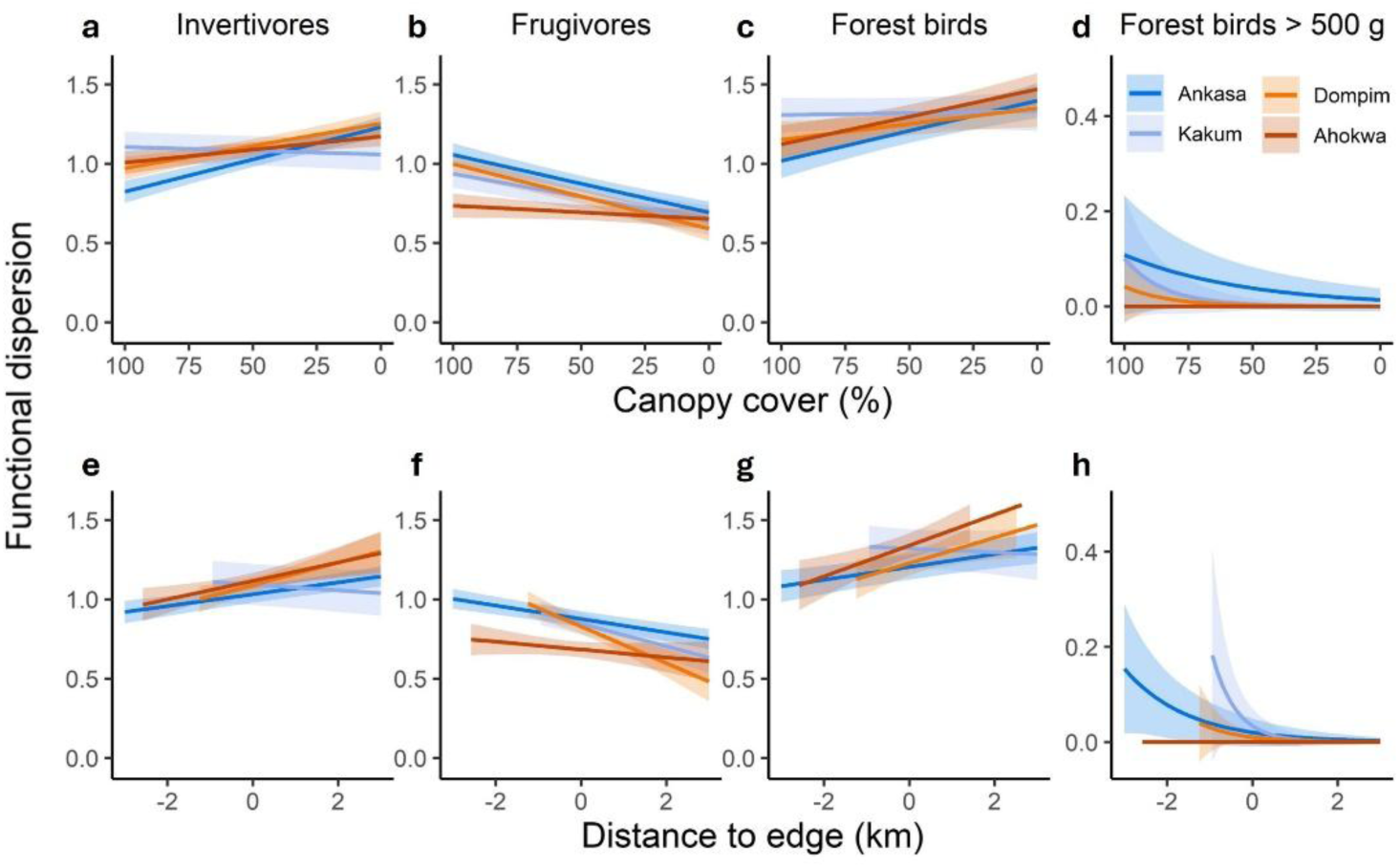
Impact of forest baseline and land-use change on the functional dispersion of bird assemblages. Results are outputs of mixed effect models examining assemblage-level functional dispersion (FDis) responses to two measures of land-use change: (**a-d**) canopy cover and (**e-h**) distance to forest edge. In **e-h**, zero indicates the forest edge, with negative values indicating distances within the forest and positive values indicating distances outside the forest. Distances to forest edge were capped at 3 km for ease of visualisation. We fitted models with bird assemblage surveys as response variable (Ankasa: *n* = 133; Kakum: *n* = 150; Dompim: *n* = 118; Ahokwa: *n* = 117). Gola was removed from analysis because the landscape surrounding the forest is dominated by agroforestry and therefore not comparable to the landscape context surrounding other study areas (see Methods). Lines indicate minimal adequate model predictions for each study site; shaded bands are 95% confidence intervals. For ease of visualisation, error bars of the insignificant slope for Ahokwa (red) were omitted in **d** and **h** and reported in SI (Table S4-S5). Note that y-axis scales vary.

**FIGURE S8.**
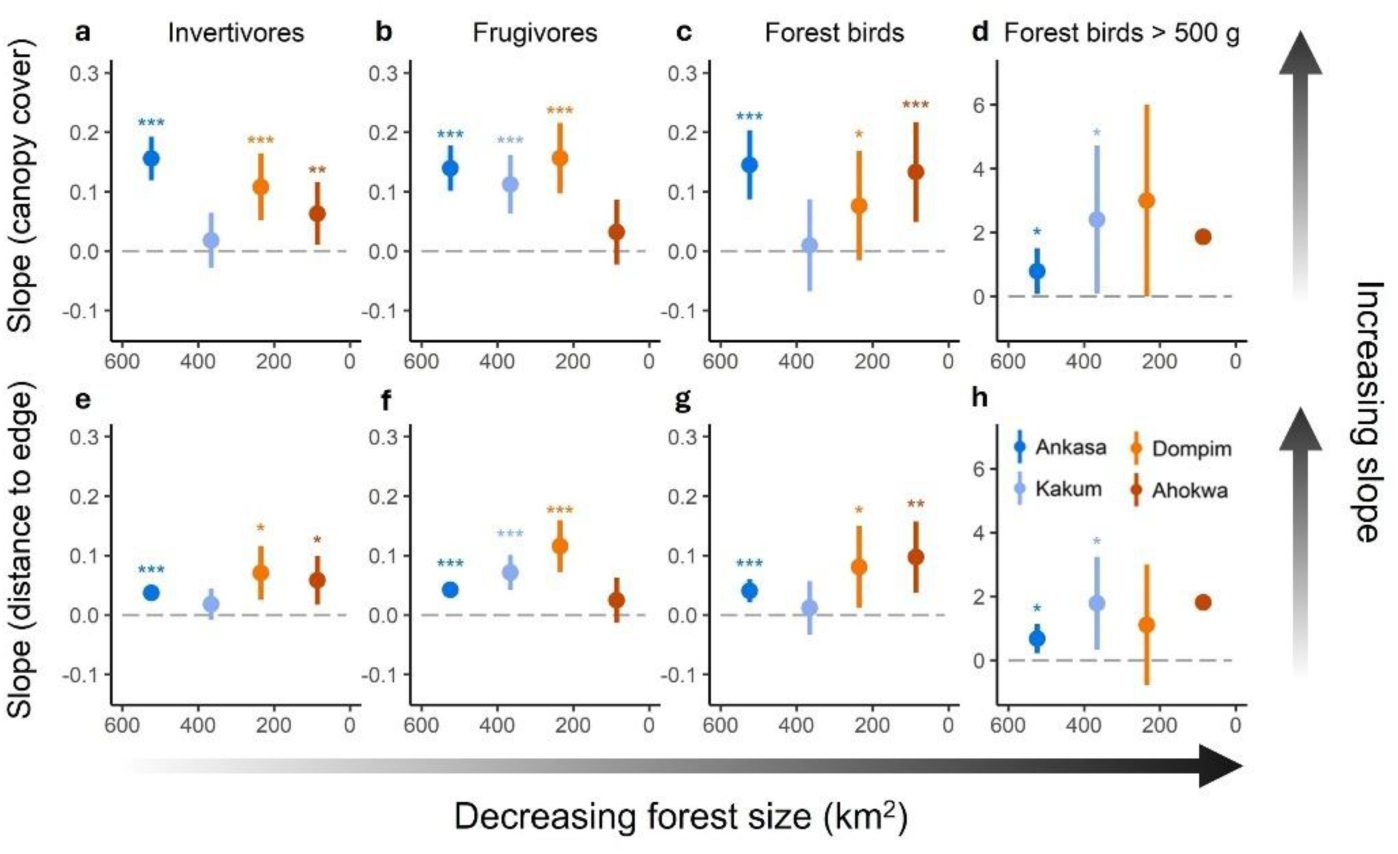
Forest size affects bird assemblage responses to land-use change. Results are outputs of mixed effect models identifying how assemblage-level functional dispersion (FDis) changes in response to two continuous measures of land-use change: canopy cover (**a-d**) and distance to forest edge (**e-h**). Results shown are coefficient estimates and 95% confidence intervals. We fitted models with assemblage surveys as a response variable (Ankasa: *n* = 133; Kakum: *n* = 150; Dompim: *n* = 118; Ahokwa: *n* = 117). Gola was removed from analysis because the landscape surrounding the forest is dominated by mixed agroforestry and therefore not comparable to the landscape context surrounding the other forests (see Methods). Slopes are absolute coefficient estimates to facilitate the comparison of steepness (i.e. absolute distance from zero) between slopes. Two contrasting patterns can be observed: large forests have steeper declines of forest-dependent species when canopy cover decreases (**a**,**b**,**d**), and small forests have steeper declines in forest-dependent species when the distance to the forest increases (**f**). For ease of visualisation, error bars of the non-significant slope for Ahokwa (red) were omitted in **d** and **h** and reported in SI (see Table S4-S5). * = p < 0.05, ** = p < 0.001, *** = p < 0.0001. Note that y-axis scales vary.

